# Genotype by environment interactions in gene regulation underlie the response to soil drying in the model grass *Brachypodium distachyon*

**DOI:** 10.1101/2024.06.17.597683

**Authors:** Jie Yun, Angela C. Burnett, Alistair Rogers, David L. Des Marais

## Abstract

Gene expression is a quantitative trait under the control of genetic and environmental factors and their interaction, so-called GxE. Understanding the mechanisms driving GxE is fundamental for ensuring stable crop performance across environments and for predicting the response of natural populations to climate change. Gene expression is regulated through complex molecular networks, yet the interactions between genotype and environment on genome-wide regulatory networks are rarely considered. In this study, we model genome-scale gene expression variation between two natural accessions of the model grass *Brachypodium distachyon* and their response to soil drying. We identified genotypic, environmental, and GxE responses in physiological, metabolic, and gene expression traits. We then identified gene regulation conservation and variation among conditions and genotypes, simplified as co-expression clusters in each combination of genotype and environmental treatment. Putative gene regulatory interactions are inferred as network edges with a graphical modelling approach, resulting in hypotheses about gene-gene interactions specific to -- or with higher affinity in -- one genotype, one treatment, or in one genotype under treatment. We further find that some gene-gene interactions are conserved across conditions such that differential expression of one gene is apparently transmitted to a target gene. These variably detected edges cluster together in co-expression modules, suggestive of different constraints or selection strength acting on specific pathways. These variable features of gene regulatory networks may represent candidates modulate environmental response via genome editing, and suggest possible targets of evolutionary change in gene regulatory networks associated with phenotypic plasticity.

## INTRODUCTION

Gene expression represents a unique class of quantitative, complex traits. Like all quantitative traits, the transcript abundance of an individual gene can take a wide range of values, can show variation in those values among samples from a population, and may be controlled by a number of intrinsic and extrinsic factors ^1–4^. And, because gene expression is a component of many quantitative traits such as morphology, life history, behavior, and physiology^5^, using tools from quantitative genetics to study gene expression may provide insight into the molecular control of these higher-order traits^6^.

Classically, variation in a quantitative trait can be modeled as a function of genetic differences among sampled individuals, of the effect of the environment on those individuals (i.e., phenotypic plasticity), and of the interaction between genotype and environment, GxE^7^. Recently, several studies have used expression QTL (eQTL) mapping to identify genetic loci that are associated with differences in the abundance of transcripts among natural accessions of plants^8–11;^ these genetic variants describe heritable, constitutive differences in expression among genotypes. Abundant studies have likewise demonstrated the strong effects that differences in environment can have on the expression of single genes or suites of genes (e.g.,^12–15)^.

Perhaps owing to the important role afforded phenotypic plasticity in plant ecology and evolution^16^, hypotheses about its molecular basis pre-date modern genomic science^17^ and remain central to studies of plant molecular biology, evolution, and biotechnology^18–24^. Molecular evolution studies or work assessing the prospects for biotechnology to enhance plant response to environmental stress often begin with transcriptomic, proteomic, or metabolomic assays of molecular responses to simulated stressors^25–29^. These -omics studies are predicated on the hypothesis that regulators of transcriptional activity offer attractive targets to manipulate suites of downstream molecular processes and, ultimately, organism-scale responses to stress (e.g., as described in ^25,30^). However, identifying genes which can be manipulated to improve performance in the field while not also incurring a fitness penalty (so-called yield drag) remains challenging ^31^. The molecular variants driving natural genetic variation in environmental response – GxE – represent a means to study how small genetic changes drive environmentally responsive physiological and developmental traits ^7^. Because such genetic variants are naturally occurring and have likely been exposed to generations of natural selection in wild populations, understanding their function *in vivo* may provide insight into how to design synthetic systems to improve stress tolerance without incurring yield drag. Genetic variants affecting GxE also represent the population-level variation on which natural selection might act to optimize fitness in response to climate change^32^ and are thus of interest for predicting the future of natural plant communities.

GxE in plant gene expression has been reported in previous studies, most often at the level of individual genes or at global scale treating each gene as an independent variable^33,34^. Such an approach can be confounded by the nature of gene regulation, in which suites of genes are often under common regulatory control ^35^; as a result, treating each gene as independent is difficult to justify statistically. Genomic features such as promoter and enhancer elements, post-transcriptional modification, and chromatin accessibility^36–40^ add further complexity to co-variance among transcript abundance *in vivo*. Collectively, the interaction between genes at the transcriptional level can be conceptualized as a gene regulation network (GRN), within which there are dependencies of expression among genes and the possible flow of information through a cell^41,42^. The common network motifs observed in GRNs imply that some regulators directly regulate or are “upstream” of other genes and may therefore exhibit pleiotropic effects when perturbed. GRNs, and the sub-networks, modules, or communities that comprise them, may be seen as an integrated molecular phenotype that allows for identifying transcriptional associations with physiological, metabolic, or developmental functions, thereby improving biological interpretability^43^.

The dependency of gene co-expression and the resulting topology of GRNs on environmental variation is poorly understood^44^. By extension, we have a limited understanding of how environmentally responding networks evolve and might be successfully manipulated using biotechnology to improve crop resilience. For example, genes exhibiting GxE in their expression might be clustered into modules, as it is hypothesized that environmental pressure results in the evolution of modules such that organismal sub-systems can function independently ^45^. A study on *Caenorhabditis elegans* found that genes exhibiting GxE in their expression have longer promoters than genome averages, with a higher density of regulatory motifs^46^. An extension of this work also proposed that GxE genes might be associated with a greater number of *trans*-acting regulators than genes showing simple genotypic differential expression^47^.

Previously, we showed that the number of genes connecting to, and the influence of genes showing GxE in expression are lower under drought stress but higher under cold stress compared to a random set of genes ^48^, again suggesting that the topology of GRNs could have been shaped by – or possibly shaped the efficacy of -- past selection in response to environmental conditions. However, that study, like most studies of environmentally dependent gene expression, assumed that co-expression relationships are invariant across genotypes or environmental treatments, which limits our ability to understand the evolution of complex gene expression phenotypes ^42,49^.

GxE is also observed in a large variety of species and its study plays important roles in fields as diverse as crop breeding and human health. However, studying GxE in plants facilitates highly replicated experiments with tightly controlled environments and genetically identical individuals. Here, we use two inbred natural accessions of the model grass *Brachypodium distachyon*^50^ to study the physiological and transcriptomic patterns of GxE in response to soil water deficit. *B. distachyon* is closely related to economically significant grasses such as wheat, barley, and oat, and it has the advantages of small stature and small genome size^50,51^. *B. distachyon* populations are found throughout the Mediterranean where they most likely exhibit a winter annual strategy – germinating in the fall or spring and setting seed before the typically hot, dry summer months^52,53^. We selected two closely related accessions, Bd21 and Bd3-1 that have similar growth rates and life histories. Moreover, the low sequence divergence in the protein coding regions of these two accessions minimize challenges with mapping RNAsequencing reads. However, we previously documented GxE in leaf water and metabolite content between Bd21 and Bd3-1^52^. The comparison between two closely related varieties is analogous to modifying a crop variety with a small number of genome edits. We use highly replicated samples of factorial combinations of genotype and environment, harnessing variation from stochastic variance of gene expression and micro-environmental heterogeneity among replicates to reconstruct regulatory interactions^54,55^. We hypothesize that environmentally responsive, genotypically specific, and genotype by environment interaction in transcript abundance and gene-gene interactions are spatially clustered in the gene regulatory network(s) of *B. distachyon*.

## RESULTS

### Physiological responses to soil drying

We first performed a drying experiment in which we measured a series of traits daily on two inbred natural accessions as soil water content was experimentally decreased from 85% of field capacity to 50% of field capacity. These measurements suggest that both accessions began reducing whole plant water usage starting on the fourth day of the dry-down (Supplementary Figure 1A, t-test adjusted by Tukey, p=0.048) and thereafter experienced a similar magnitude of soil drying stress with most assayed traits exhibiting clear differences between control and drying treatments by Day 6. Linear mixed models show that both accessions respond similarly along this dry-down, with the exception of leaf glucose and fructose contents, which showed significant increase starting on Day 5 in Bd3-1 (z-test, p<0.0001) and Day 6 in Bd21 (p<0.0001; Supplementary Figure 1H and Supplementary Figure 2B).

We next performed a more highly replicated experiment (n=48-58 for each accession in each treatment) focused on contrasts between the two accessions at soil water contents of 85% and 55%, corresponding to Day 6 of the dry-down process. These plants constitute the sampling for all following analyses.

Leaf metabolite content was affected by soil drying. Leaf starch content decreased on Day 6 (Figure 1G; Chisq test and p=1.6e-10). Sucrose shows no treatment effects in the leaf (Figure 1E; p=0.48). The degradation of starch and the catabolism of sucrose might contribute to the accumulation of glucose and fructose, both of which are strongly elevated in our Day 6 samples (Figure 1C, D; both p<2.2e-16). However, leaf glucose increases to a larger degree in Bd3-1 than in Bd21, shown as a significant GxE response (Figure 1C, p=0.0015). Another important group of carbohydrates in cool-season Pooid grasses like *Brachypodium* are fructans, which may serve as a short-term storage carbohydrate when sucrose exceeds demand ^56^. Here, we assayed high-degree polymerized fructan, which showed detectable decreases in leaves on Day 6 (Figure 1H; p=1.6e-7), similar to what we observed for starch. Low-degree polymerized fructan also decrease slightly (Figure 1F; p= 0.055), similar to sucrose. In general, most carbohydrates show a genotypic effect with higher per-dry-mass values in Bd3-1. Amino acids per unit leaf mass was higher in droughted plants on Day 6 (Figure 1J; p<2.2e-16). Total protein content increased in Bd3-1 and marginally increased in Bd21 (Figure 1I; p=0.00015), showing a GxE response (p=0.034).

**Figure 1.**
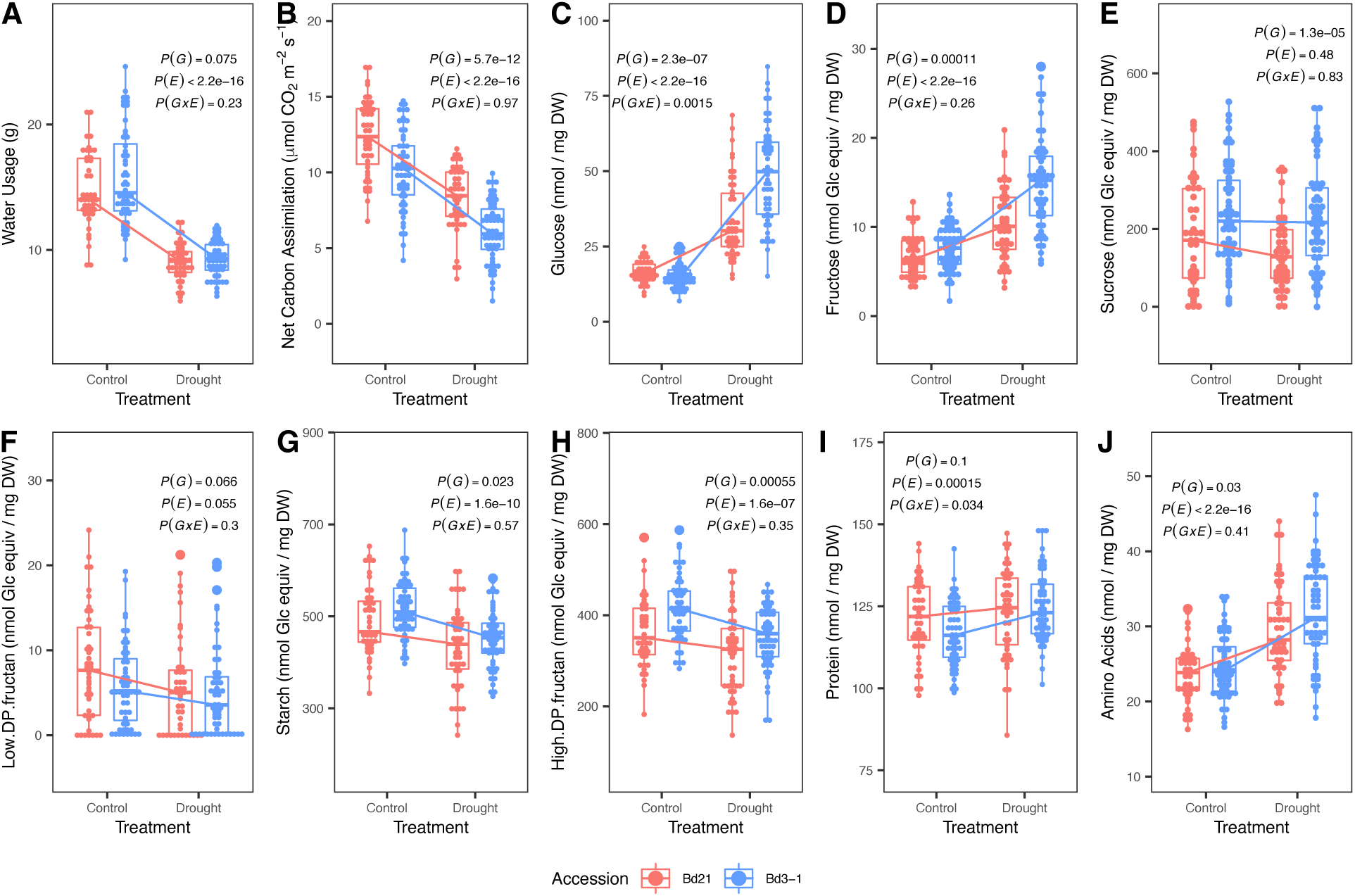
Leaf blade traits of Bd21 and Bd3-1 on Day 6. The p-values of effect terms were determined using (generalized) linear (mixed) models, by removing each term and performing likelihood ratio test (anova and Chisq test, with maximum likelihood mode if models have random factors). DW = Dry Weight of tissue. Glc Equiv = glucose equivalents. N = 48-58 for each combination of accession and treatment.

### Factors affecting the transcript abundance of individual genes

The transcripts of 29,740 gene models were identified in our RNASequencing data, representing 91.4% of those in the Bd21 v3.0 reference genome. Principle component analysis shows clear grouping of samples by library type (Figure 2A), in which the first PC separates the two genotypes and explains 15.5% of total sample variance and the second PC separates the two treatments and explains 6.4% of total variance. 26,388 genes passed our filtering criteria and are included in the following analysis of by-gene differential expression (Supplementary Data Set 1). At an FDR=0.05, 4725 genes are constitutively higher, and 4087 genes are constitutively lower in Bd21 as compared to Bd3-1 (i.e., show a significant genotypic effect). 3111 genes are constitutively higher, and 2832 genes are lower in drought than control (environmental effect); among these, 3066 genes exhibit both significant genotype and environment effects. In Bd21, 4024 and 3983 genes are up- or down-regulated, respectively, under drought. In Bd3-1, 3997 and 4010 genes are up- or down-regulated, respectively, under drought (Figure 2B). 538 genes exhibit significant GxE interaction at FDR = 0.1 (Figure 2B; hereafter “GxE genes”). Among these GxE genes, 107 are up-regulated and 66 are down-regulated in both accessions but with differing magnitudes of response in the two accessions (Figure 2B). 135 genes are uniquely up-regulated and 99 uniquely down-regulated in response to drying in Bd3-1 (so-called “conditional neutrality;”^7^); similarly, 39 genes are uniquely up -regulated and 60 uniquely down-regulated in Bd21. Only 21 genes are expressed in different directions in response to the drought treatment in the two accessions.

**Figure 2.**
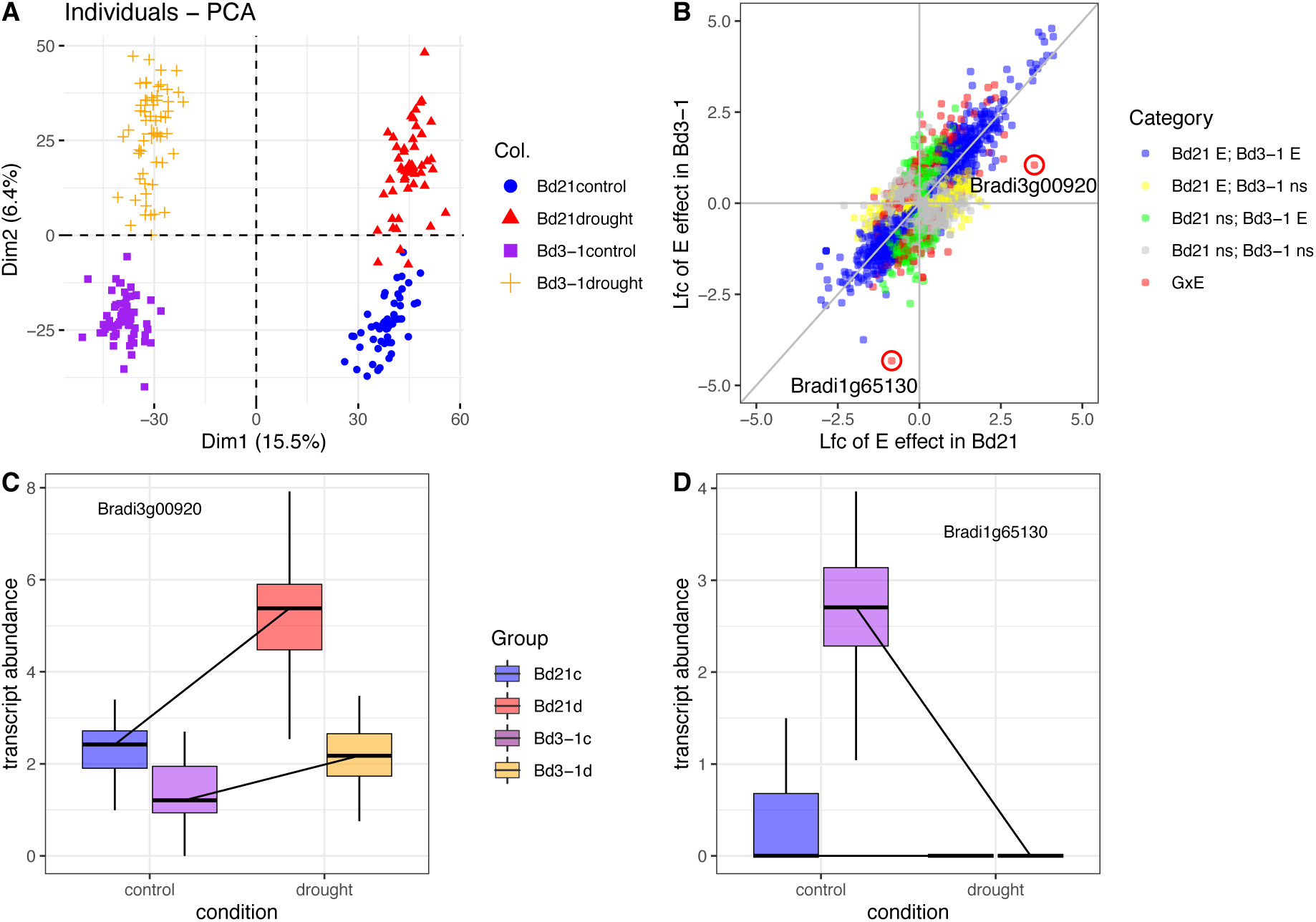
Gene expression diversity. **A)** PCA shows clustering of RNASequencing library by accession and environmental treatment. **B)** Factors affecting the transcript abundance of individual genes. Each point represents a gene model in the *B. distachyon* Bd21 genome. Lfc = Log-fold change of transcript abundance in response to soil drying. Colors indicate the significance of treatment effect on Bd21 and Bd3-1 samples; genes with a significant GxE effect are highlighted in red and the two genes plotted in **C and D** are circled. Boxplots show two genes with large GxE effect: **C)** Bradi3g00920 and **D)** Bradi1g65130, gene expression reaction norm based on log2(norm_count+1).

There are 40 annotated transcription factors among the 538 GxE genes, including a predicted MYB-related transcription factor with the largest GxE effect (−1.5 log-fold change). GO terms enriched among GxE genes include processes related to chlorophyll, starch and organic substances, amino acids, translation, and protein folding processes, and response to stimulus (Supplementary Data Set 2).

### Variation in gene-gene interaction within co-expression modules

We next test the hypothesis that groups of putatively co-regulated genes show genotypic, environmental, or GxE effects. The high replication used in our RNA sequencing experiment (N = 48-58, subsampled to 48 samples for each library type) affords us the power to estimate co-expression relationships independently in each of the four library types (Bd21 control watering, Bd21 drought, Bd3-1 control, Bd3-1 drought) using Weighted Gene Co-expression Network Analysis (WGCNA)^57^. Genes in a module might represent a group of genes commonly activated by regulatory factors and sharing cellular functions^43^. Depending on the two parameter settings implemented in WGCNA we identified 14-24 modules in Bd21 control, 14-18 modules in Bd21 drought, 14-16 modules in Bd3-1 control, and 15-20 modules in Bd3-1 drought (Figure 3).

**Figure 3.**
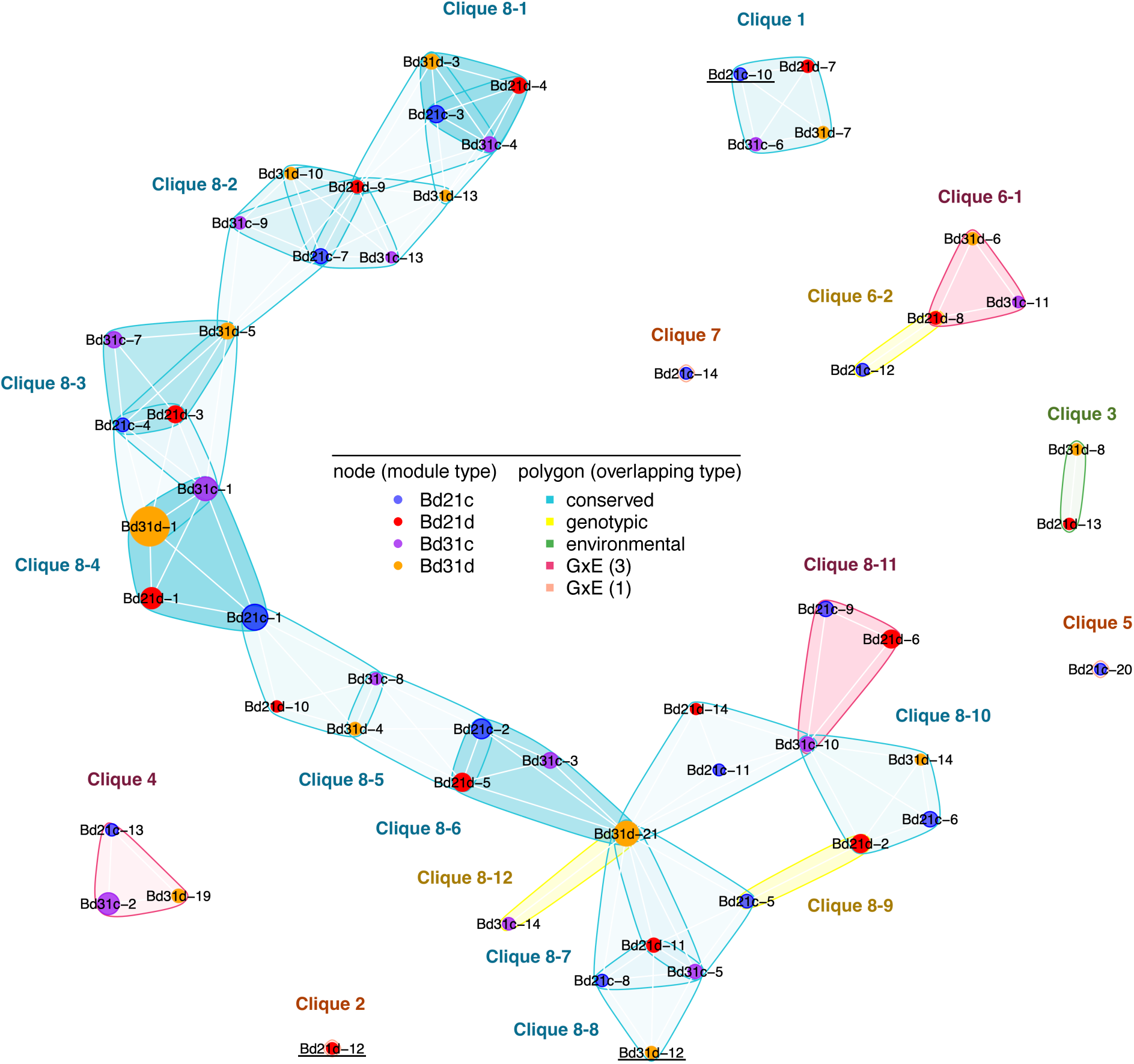
Conservation of gene co-expression modules among genotypes and treatments. Nodes represent modules identified in each of the four library types, with the size of the node corresponding to the number of genes in each module (red, Bd21d; blue, Bd21c; orange, Bd3-1d; purple, Bd3-1c). Clusters of modules enclosed by a polygon -- cliques -- show significant gene overlap using the high-dimensional Fisher’s Exact Test (Data are available in Supplementary Data Set 5); polygon color denotes the pattern of module conservation among libraries (teal, four-module stable overlap; pink, three-module overlap; green, two-module of same environment overlap; yellow, two-module of same genotype overlap; and orange, module uniquely identified in a single library type) with the transparency of the color proportional to the number of genes in common. Underlined module names are shown as inferred network in Figure 4.

We first identify modules that overlap among the four library types using a high dimensional Fisher’s exact test^58^, testing the hypothesis that a set of modules identified within or between library types share significantly more genes than expected. We interrogated these “cliques” of overlapping modules for four general patterns. In the first pattern, we reason that a clique comprised of one module from each of the four library types may represent gene co-expression relationships that are conserved across genotype and treatment (Figure 3, teal polygons). We find 40 such modules comprising 20 overlapping cliques, among which 36 modules show significant GO enrichments.

Clique 1 is comprised of four modules that have common GO terms of circadian rhythm. Clique 8 is comprised of a chain of nine partially overlapping cliques enriched for core cellular processes following a spatial sequence, including electron transport chain, cellular respiration, protein translation, photosynthesis light reaction, and glycogen metabolic process (Table 1). Some modules in conserved cliques are highly correlated with physiological traits. In particular, Bd3-1d21 in Clique 8-6 is correlated with daily water usage (p-value=4.4e-5) while Bd3-1d4 in Clique 8-5 is correlated with both daily water usage (p-value=1.5e-4), and tissue water content (p-value=1.9e-4). Both modules are enriched with GO terms associated with photosynthesis. Bd3-1d12 in Clique 8-8 is correlated with leaf glucose content (p-value=6.4e-5) and partially overlaps with Clique 8-7 which is enriched for glycogen metabolic process.

**Table 1.** Common GO terms at lowest level among modules in overlapping groups (based on all module versions FDR=0.1) and correlated traits (based on moderate size module version, in specific library type, only p<0.001 are reported, not corrected by multiple test). DGE enrichment in modules are shown (based on constraint module size version, only p<0.001 are reported, not corrected by multiple test, and the percentage is shown if > 75%). Cliques with no common GO terms or correlated traits identified are not shown. Complete lists are available in Supplementary Data Set 6-8.

| Name (# of included modules) | Common GO | Correlated Trait (p<1e-03) | DGE Enrichment (p<1e-03; percentage when >75%) |
| --- | --- | --- | --- |
| Clique 1 (4) | circadian rhythm |  | Bd3-1c6: E |
| Clique 2 (1) |  | glucose (1.1e-05) | E (97%),GxE |
| Clique 3 (2) |  | Bd21d13: Glucose (9.4e-08)<br>Bd3-1d8: Glucose (1.6e-06) | Bd21d13: E (100%);<br>Bd3-1d8: E (92%),GxE |
| Clique 4 (3) |  | Bd3-1d19: leaf water content (8.5e-04) | Bd21c13: G (76%);<br>Bd3-1c2: E,G,GxE;<br>Bd3-1d19: G (97%) |
| Clique 5 (1) | phosphatidylethanolamine biosynthetic process |  |  |
| Clique 7 (1) | glutathione metabolic process; alternative respiration; response to chemical |  | GxE |
| Clique 8-1 (4) | ribosomal small subunit assembly; translation; photosynthetic electron transport in photosystem II; photorespiration; ATP synthesis coupled proton transport; reductive pentose-phosphate cycle; electron transport coupled proton transport, respiratory electron transport chain |  |  |
| Clique 8-2 (4) | electron transport chain, cellular nitrogen compound biosynthetic process, cellular respiration |  |  |
| Clique 8-3 (4) | translation |  |  |
| Clique 8-4 (4) | cytoplasmic translation; regulation of actin polymerization or depolymerization; ribosomal large subunit biogenesis, protein maturation |  |  |
| Clique 8-5 (4) | Photorespiration; reductive pentose-phosphate cycle | Bd3-1d4: Leaf water content (1.5e-04) daily water usage (1.9e-04) | Bd21c2: E,G; Bd21d5: E (82%); |
| Clique 8-6 (4) | response to light stimulus; photosynthesis, light harvesting in photosystem I; photosystem II stabilization; pigment biosynthetic process | Bd3-1d21: daily water usage (4.4e-05) | Bd21c2: E,G; Bd21d5: E (82%);<br>Bd3-1c3: E; Bd3-1d21: E,G,GxE |
| Clique 8-7 (4) | glycogen metabolic process |  | Bd21c8: E,G; Bd21d11: E,G,GxE;<br>Bd3-1c5: E,G,GxE;<br>Bd3-1d21: E,G,GxE |
| Clique 8-8 (4) |  | Bd3-1d12: Glucose (6.4e-05) | Bd21c8: E,G; Bd21d11: E,G,GxE;<br>Bd3-1c5: E,G,GxE;<br>Bd3-1d12: E,GxE |
| Clique 8-9 (2) | response to wounding; phenylpropanoid biosynthetic process; regulation of defense response; monocarboxylic acid metabolic process |  | Bd21c5: G,GxE;<br>Bd21d2: E,G,GxE |
| Clique 8-11 (3) | plant-type primary cell wall biogenesis; plant-type secondary cell wall biogenesis; cellulose biosynthetic process |  | Bd21c9: E,G (100%);<br>Bd21d6: E,G (95%);<br>Bd3-1c10: E,G (82%) |
| Clique 8-12 (2) | terpenoid biosynthetic process; chlorophyll metabolic process; pigment metabolic process | Bd3-1d21: daily water usage (4.4e-05) | Bd3-1c14: E,G (76%)<br>Bd3-1d21: E,G,GxE |

The second pattern is a clique comprised of two modules from the same genotype and both treatments (Figure 3, yellow polygons). We find three such cliques, which may represent genes that are only co-regulated in one genotype. For example, Clique 8-9 solely comprises modules in Bd21 and is enriched with functions such as response to wounding and phenylpropanoid biosynthetic processes (including *B. distachyon* orthologs of genes encoding C4H (Bradi2g31510), PAL (Bradi5g15830), 4CL (Bradi3g52350) and F3H (Bradi2g54090, Bradi5g19250, Bradi2g54090), F3’H(Bradi4g31380), PPO (Bradi2g52260) and an R2R3 MYB transcription factor (Bradi2g11676) which regulates phenylpropanoid biosynthesis in other species; Supplementary Data Sets 3-4). Clique 8-12 is comprised solely of modules in Bd3-1 and is enriched in chlorophyll and pigment metabolic processes and terpenoid biosynthetic processes; it contains five NAC regulators/domain related genes (Bradi1g17480, Bradi1g63600, Bradi3g59380, Bradi4g02060, Bradi4g44000) as well as a chlorophyll A-B binding protein (Bradi4g31257) and a putative senescence-inducible chloroplast stay-green protein (Bradi4g36060).

The third type of clique is treatment specific, composed of two overlapping modules from different genotypes but the same environmental condition (Figure 3, green polygons). We identified one such case, Clique 3. Both modules in this clique are correlated with leaf glucose content (p-value=9.4e-8 and 1.6e-6 for Bd21 and Bd3-1, respectively). Although the two modules have no GO terms in common, Bd21d13 is enriched with GO terms including response to water and contains two RAB18 orthologs (Bradi1g37410, Bradi3g43870), one rice ASR5 ortholog (Bradi4g24650), and one IAR4 (Bradi1g44480). As expected, many genes in Bd21d13 -- including the two RAB18 genes -- have very low expression in the control condition.

Finally, a clique may be comprised of modules either from a single library type or from three library types and excluding the fourth library type; we interpret this pattern to represent modules exhibiting GxE (Figure 3, pink or orange polygons). There are three cliques which are unique to a single module in a single library type. Among these, Clique 2 is comprised solely by a module identified in Bd21 drought and, although not enriched for any GO terms, the module is highly correlated with leaf glucose content (p-value=1.1e-5). The annotations of several genes in this module relate to abscisic acid (ABA) signaling and regulation. Moreover, the proximal promoters of genes in this module are enriched with a predicted bZIP motif (p-value = 2.6e-7). Clique 5 comprises only one module from Bd21c and is enriched with GO term of phosphatidylethanolamine biosynthesis process. Clique 7 is identified in Bd21 control watering, enriched with GO terms of glutathione metabolic process and response to chemical. We also identified three cliques comprised of modules from three library types with the fourth condition unrepresented. Among these, Clique 8-11 lacks a Bd3-1d module and the three included modules have GO terms in common related to cell wall biogenesis. Many of the genes in Clique 8-11’s modules are not included in any reconstructed module in Bd3-1d.

### Inferring regulatory interactions within modules

The preceding analyses demonstrate that the gene compositions of some co-expression modules are conserved across genotypes and treatments while others vary as a function of genotype, treatment, or GxE. Because co-expression may arise from regulatory interactions, we next assess a possible cause of module-level variation: variation in regulatory interactions among genes in modules. We use graphical models to infer the conditional independence of genes to estimate causal relationships among them, allowing us to infer the directionality of gene regulation. Here we work with the simplifying assumption that the regulator(s) of a focal gene is in the same module as that gene. We focus on three exemplar modules. Since the regulation is based on estimates of transcript abundance, gene “interaction” might represent either direct transcription factor-target regulation or indirect regulation that might arise from system-level phenomena such as metabolism or the cell cycle ^59^.

Bd21d12 is the sole module in Clique 2, which we above infer to be a module whose coherence is GxE-dependent. Many of the genes in Bd21d12 are annotated with functions in ABA biosynthesis and signaling and two of these genes are predicted to act upstream many other genes in the module (Figure 4A red fill): Bradi1g13760, an ortholog of AtNCED2, a 9-cis-epoxycarotenoid dioxygenase, and Bradi3g52660, orthologous to AtCYP707A1, with inferred ABA 8’-hydroxylase activity. The module also contains genes whose orthologs are ABA-induced or ABA-dependent regulators (Figure 4A orange fill), including Bradi4g39520 (orthologous to a transcription repressor, AtAITR3), Bradi3g50220 (orthologous to ABA-responsive ATHB-7), Bradi2g15940 (orthologous to AtGBF2, an ABA regulated bZIP transcription factor), and Bradi2g41950 and Bradi2g18510 (orthologous to AtHAB2 and AtHAB1, respectively). Other genes are associated with drought response in Arabidopsis (Figure 4A blue fill), including stomatal movement (Bradi2g20660, Bradi3g39800), wax biosynthesis (Bradi2g23740), and a late embryo abundant (LEA) protein (Bradi1g51770). The gene with strongest GxE effect in our dataset, Bradi3g00920 (Figure 2C), has the highest eigenvalue in this module.

**Figure 4.**
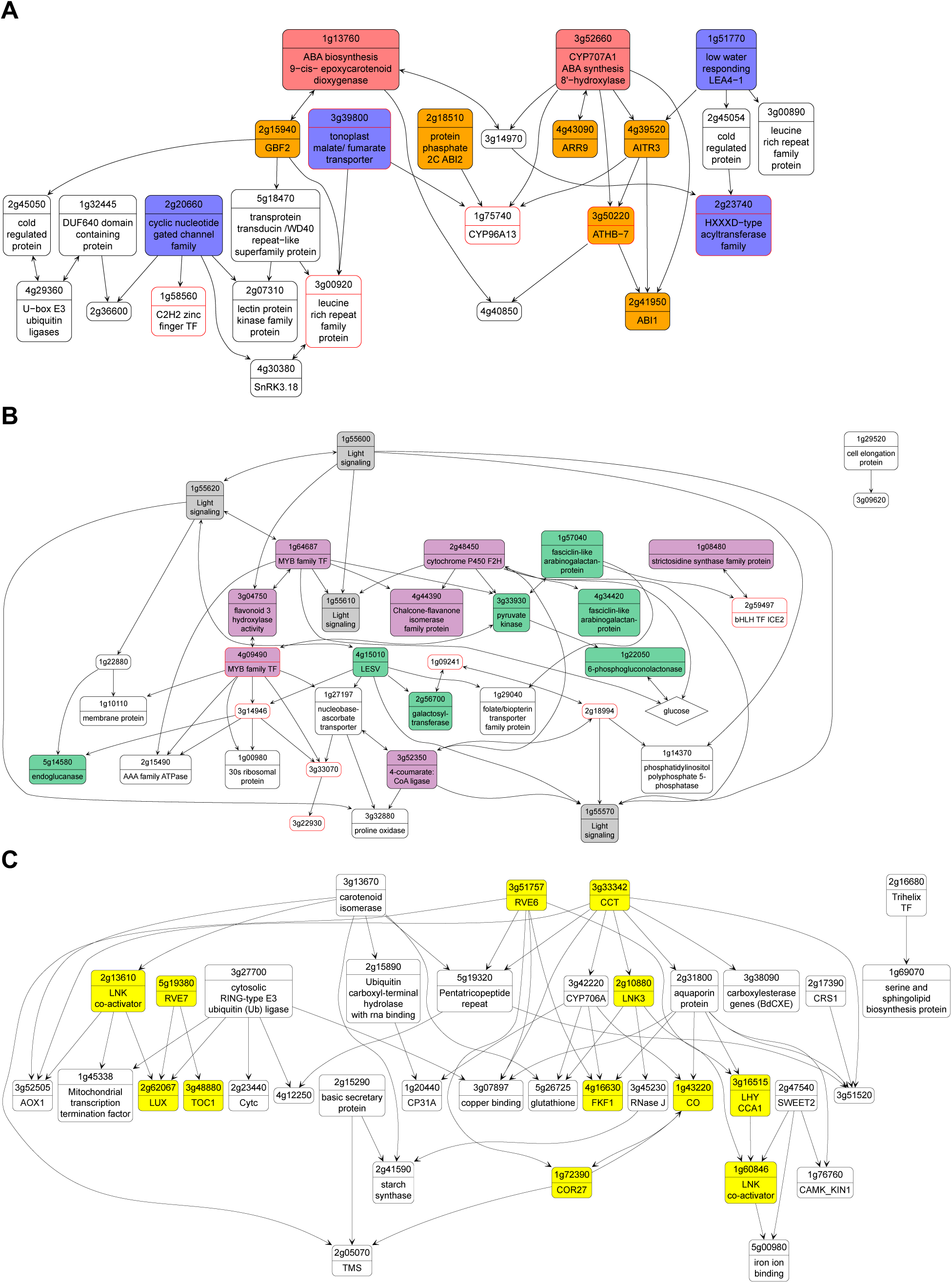
Inferred directed interactions within modules. **A)** Module 12 in Bd21d, **B)** Module 14 in Bd3-1d and **C)** Module 10 in Bd21d inferred using the UTI-GSP method. Nodes are genes and edges are inferred causal interactions. Edges with two arrows indicate either the direction of interaction is uncertain or that the interaction is inferred to be in both directions. Second row of each node shows annotation of orthologs in Arabidopsis or rice. Glucose is included as a node in module Bd3-1d12 as an example of a module with a significant correlation between leaf glucose content and the module eigengene (Table 1). Genes exhibiting a significant pattern of GxE in our DGE analyses are highlighted with red outline.

Module 12 in Bd3-1d is correlated with leaf glucose content and is a member of Clique 8-8 (Figure 4B). Bradi1g55600 is inferred to be upstream of many other genes in Bd3-1d12 (Figure 4B grey fill); its Arabidopsis ortholog is an early light-induced protein (ELIP2) and regulates chlorophyll synthesis^60^. Paralogs of Bradi1g55600 are also present in this module. Downstream in the inferred network there are genes whose Arabidopsis orthologs encode enzymes in primary metabolism (Figure 4B green fill), including Bradi3g33930 (pyruvate kinase), Bradi1g22050 (6-phosphogluconolactonase in the oxidative phosphorylation pathway), Bradi5g14580 (an endoglucanase), Bradi2g56700 (a galactosyltransferase), and Bradi1g18650 (a 4-alpha-glucanotransferase in starch degradation). Above, we reported that leaf glucose content is correlated with this module; when we include glucose as a node in the module, we find that it is predicted to directly interact with Bradi1g22050, the 6-phosphogluconolactonase. Module Bd3-1d12 also contains Bradi4g15010, whose Arabidopsis ortholog is predicted to have starch synthesizing and degrading activity (LESV). Additional genes in this module have predicted functions in secondary metabolism (Figure 4B purple fill), including orthologs of an Arabidopsis MYB family transcription factor involved in flavonol biosynthesis (Bradi1g64687), flavonoid 3’-hydroxylase (Bradi3g04750), flavanone 2-hydroxylase (Bradi2g48450) and a 4-coumarate: CoA ligase in the phenylpropanoid pathway (Bradi3g52350). Notably, several genes with unknown functions are connected in this module and exhibit GxE in their individual transcript abundances.

Finally, we report an example of a clique that is conserved in all four library types. We investigated its constituent module, Bd21d10, as a representative of the clique and find that it is enriched in genes relate to circadian processes (Figure 4C, yellow fill). There are five predicted MYB TFs in Bd21d10, including Bradi3g51757 (orthologous to AtRVE6), Bradi2g62067 (orthologous to AtBOA and OsLUX), Bradi5g19380 and Bradi1g29680 (orthologous to AtRVE2/7), Bradi3g16515 (orthologous to AtLHY1, and OsCCA1). Additional genes in this module are orthologous to CONSTANS (CO) and related proteins in Arabidopsis. Genes in circadian-controlled pathways show conserved oscillation over 24 hour intervals^61^, suggesting that stochastic clock variance among our sampled individuals may have provided the signal for reconstructing these conserved interactions in our data.

### Genes predicted to interact exhibit various patterns of differential gene expression

Several of the reconstructed co-expression modules are enriched with genes that exhibit differential gene expression (Table 1). We next ask whether pairs of genes predicted to interact in our directed regulatory networks (Figure 4) share a common pattern of differential gene expression. For example, two genes sharing a common edge may differ in their response to genotype (e.g., one gene shows no DGE and the second gene shows a genotypic effect on gene expression), environment, GxE, or no difference. (See Supplementary File 1 for illustrative examples.) We summarize these different classes of shared or divergent DGE in Supplementary Table 1 and Supplementary Figure 3. For example, of 41 total edges in module Bd21d12 (which we only observe in Bd21 under drought), there are five edges showing G changes, 11 edges showing E changes, 12 edges showing both G and E changes, 11 edges showing GxE changes, and two edges linking nodes with no difference in DGE. Of 62 total edges in Bd21c5 (unique to Bd21), there are 23 edges showing no DGE changes, 25 edges showing G changes, nine edges showing E changes, eight edges showing both G and E changes, and no edges showing GxE changes.

### Genetic and environmental effects on the interactions among genes

An edge between nodes in our inferred gene regulation network represents a hypothesis about the regulatory interaction between two genes. The high biological replication of our RNASequencing across four library types enables us to fit linear models to explore the effects of genotype, environment, and their interaction on the expression relationship between pairs of genes. There are four patterns of interest in our dataset.

First, the regulatory interaction between two genes may be present under one or some combinations of genotype and treatment but absent in others; this pattern would be diagnosed by identifying a significant change of the slope of the regression relationship between two genes. Environmentally dependent regulatory interaction is one possible cause of such a pattern (Figure 5B), and we see many clear examples of environmental regulation. In module Bd21d12, described above, the regression of Bradi4g40850 (unknown function) on upstream genes shows a significant E effect of slope from gene Bradi3g50220 (orthologous to the ABA-dependent ATHB-7 transcription factor; Figure 5B). Similarly, in drought specific module Bd21d13, the regression of Bradi1g37410 (which encodes a dehydrin family protein) on upstream genes suggests that Bradi1g44480 (predicted dehydrogenase E1 component domain containing protein) regulates Bradi1g37410 only in drought (Supplementary Figure 4A). Across modules, we see clustering of such context-dependent edges, such as module Bd21d13 in Clique 3 – reconstructed only under drought conditions – comprising 32 edges of which 14 (44%) are environmentally dependent (Supplementary Table 1).

**Figure 5.**
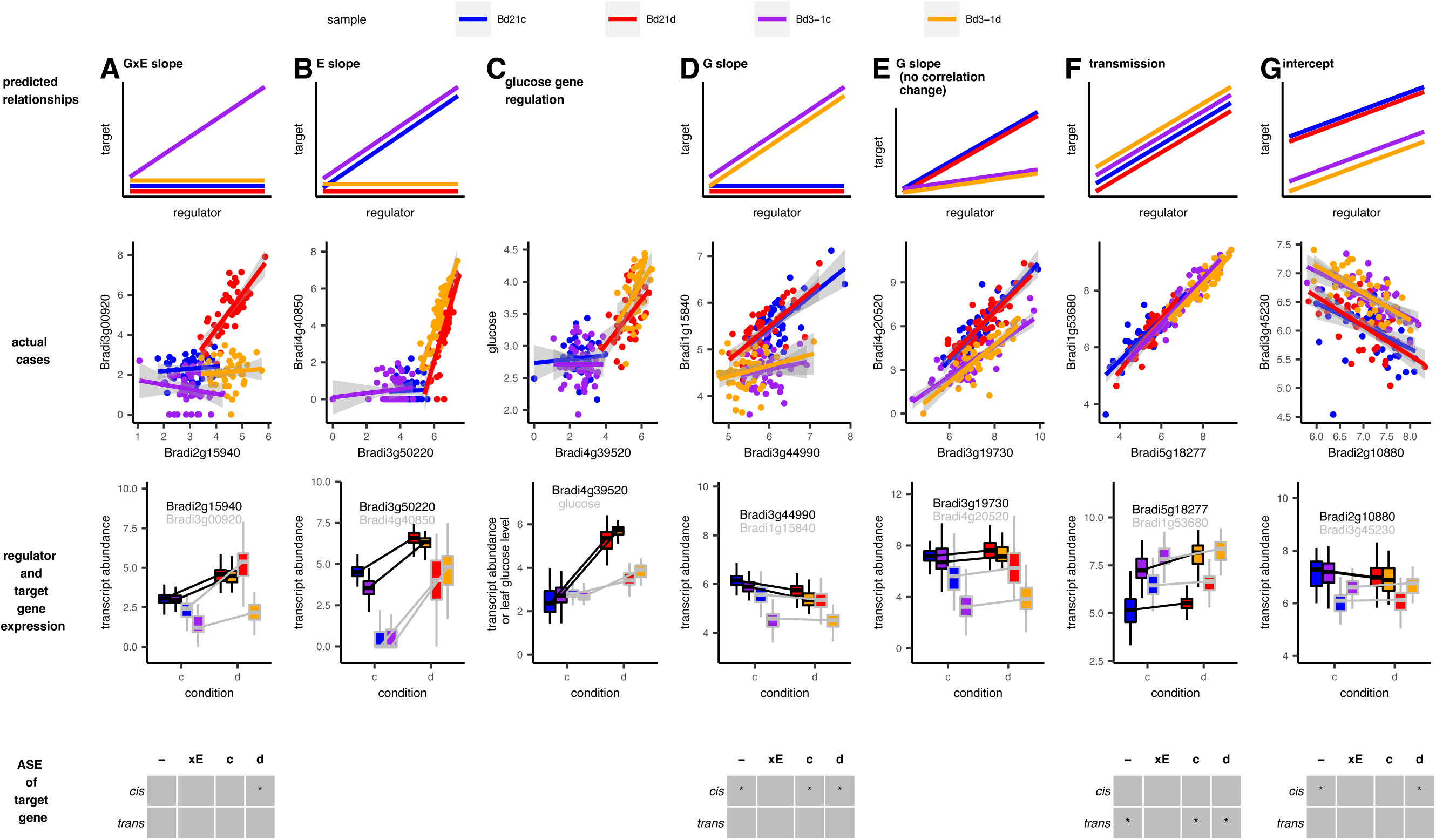
Predicted and actual examples of regulations between pairs of genes selected. **A-B, D-G)** show regulation slope changes, transmission, and regulation intercept changes. These are illustrated with linear regression between regulator and target, the expression level of each gene, and allele specific expression (ASE) types. ASE types include constitutive regulation (-) and environment specific (xE) detected in full model (see method), or regulation detected in control (c) and drought (d) condition. P-value <0.1 and not corrected by multiple tests. **C)** regression of log-transformed leaf glucose content on upstream identified gene Bradi4g39520 as a putative regulator of leaf glucose content. Transcript abundances are log2 (norm_count+1).

A regulatory interaction between two genes also may be observed in only one of the two genotypes (Figure 5D). For example, in Bd21 specific module Bd21c5, the regression of Bradi1g15840 (putative allene oxide cyclase in jasmonic acid synthesis) on upstream genes suggest Bradi3g44990 (putative cell-wall invertase) regulates Bradi1g15840 only in Bd21 (Figure 5D). Consistent with the observation that the genes in Bd21c5 are only co-expressed in Bd21, we find that Bd21c5 has 12 edges showing a significant effect of genotype among its among 65 edges (18%; Supplementary Table 1).

We might also observe that a regression is only significant in a single library type, indicative of GxE for a given pairwise regulatory interaction (Figure 5A). In module Bd21d12, discussed above and enriched for ABA-responsive genes, the regression of Bradi3g00920 (Leucine Rich Repeat family protein) on upstream genes suggests that Bradi2g15940 (GBF2) interacts with Bradi3g00920 with a significant GxE effect of the slope (Figure 5A).

In the second general pattern, we observe a change in the slope of the regression of the target gene on its putative regulator without a complete loss of the correlation; this might be caused if the binding interaction between a transcription factor and a regulatory motif is stronger in one genotype than a second genotype (Figure 5E). As one example, Bradi4g20520 (unknown function) is predicted to regulate Bradi3g19730 (jacalin-like lectin domain containing protein) in all library types but with a greater slope in the Bd21 libraries than in Bd3-1 libraries.

Third, the baseline expression of genes might be different across library types (Figure 5G), possibly due to the presence of cofactors or the general resource and energy availability of the cell; this type of regulation may be detected as a difference in the regression line’s intercept among library types. In module Bd21c10, the Bradi3g45230 (RNA-metabolising metallo-beta-lactamase) regression on Bradi2g10880 (LNK3) shows is consistent with the hypothesis that their interaction is similar in all library types, but that the baseline expression in Bd3-1 is constitutively higher than in Bd21. This pattern is observed as a higher intercept in Bd3-1 libraries which leads to genetically dependent expression of the downstream gene. Such intercept differences as a function of environment, genotype, or GxE are widely found among the pairwise interactions identified in our data (Supplementary Table 1).

The final pattern is no difference in the pairwise regulation between genes among library types but an apparent transmission of a regulatory response between the genes. This pattern is manifested as neither a significant slope change nor an intercept effect in the regression, and a common differential expression between putative regulator and its target gene (e.g. each gene exhibits a significant effect of GxE on its own transcript abundance). One such case is Bradi5g18277 (predicted serine-type endopeptidase) and Bradi1g53680 (predicted zinc transporter 3 precursor) in module Bd31c16 (Figure 5F). In module Bd31d12, GxE genes are found more likely to be regulated by genes exhibiting GxE than by non-GxE genes in the module (t-test, p-value = 0.013; see Supplementary Figure 4B for an example).

Finally, we use our regression approach to study the transcriptional control of leaf glucose content, which shows a significant GxE effect in our metabolic assays (Figure 1H) and is highly correlated with four module eigengenes (module Bd21d12, Bd21d13, Bd3-1d8, Bd3-1d12; Table 1). Nine genes are inferred to regulate glucose when it is included as a node in these modules. Among these, we find that Bradi4g39520 only exhibits an interaction with glucose under drought stress (Figure 5C); the glucose content is same between the genotypes and is not dependent on the expression level of Bradi4g39520 under control conditions. Bradi4g39520 is an ortholog of AITR3, an ABA-induced transcription repressor that acts as feedback regulator in ABA signaling in Arabidopsis.

### Identifying allele-specific expression in target genes

We next used allele specific expression assays to infer whether the observed genetic or GxE-associated changes in regulatory interactions arise from genetic factors acting in *cis* or in *trans*. In the above example of genotype-dependent interaction between Bradi3g44990 and Bradi1g15840 (Figure 5D), Bradi1g15840 is inferred to be *cis-* regulated (Supplementary Figure 5B). In the example of putative GxE regulation of Bradi3g00920 by Bradi2g15940 (Figure 5A), we identified Bradi3g00920 is regulated by a *cis*-acting factor under drought stress (Supplementary Figure 5A). In the example of an apparent transmission of expression GxE from Bradi5g18277 to Bradi1g53680 (Figure 5F), we observed that transcript variation of Bradi1g53680 is controlled by a *trans*-acting variant (Supplementary Figure 5C). We observe that the genotype-dependent regression intercept of Bradi3g45230 is associated with a cis-variant (Figure 5G and Supplementary Figure 5D), consistent with a model in which Bradi2g10880 regulates Bradi3g45230 differently in two genotypes due to a variable enhancer or other *cis-* regulatory element.

## DISCUSSION

Gene expression has long been a focus of both evolutionary and functional genetic studies. Here, we aimed to integrate these two perspectives by studying the environmental and genetic control of regulatory interactions between genes using highly replicated RNASequencing libraries of *Brachypodium distachyon* accessions exposed to moderate soil drying and control conditions.

We extend past work showing that the expression of individual genes can vary as a function of genotype and environmental treatment to demonstrate variation in interactions between genes. One example highlighted throughout the analyses in this study is Bradi3g00920, which shows a pattern of GxE in its transcript abundance. By developing a directed graphical network for the module in which the gene is located, we infer that Bradi3g00920 is relatively downstream of several other genes putatively regulating it. Multiple regression of its gene expression suggests that Bradi3g00920 is regulated by Bradi2g19540 only under drought in Bd21. We infer that this pattern suggests a possible cause of the observed GxE in gene expression for Bradi3g00920. Further, allele-specific expression reveals that this gene is associated with a *cis-*regulated expression pattern.

We next consider how a differentially expressed gene may transmit its expression pattern through its interactions with other genes. Here, we refer to the correlated pattern of differential expression as “transmission.” As shown in the example in Figure 6F, an inferred target gene (Bradi1g53680) shows the same dependency on a putative regulator (Bradi5g18277) across all library types. Indeed, the target gene is found to be *trans*-regulated, suggesting that the genetic variant(s) causing the differential expression is in the direct regulator or upstream of it. An implication of this finding is that many *trans*-acting loci in QTL studies might include such gene expression transmission and not a protein coding change in the *trans-*acting regulator. The transmission of the expression response is also a direct example showing certain genetic variants can affect more than one gene, and that the expression of individual genes should not be considered independent variables in transcriptome studies.

A second key implication of our work is that variation in gene-gene interactions observed in modules can lead to a module not being reconstructed in certain combinations of genotype and environment. For example, 44% of the inferred edges in module Bd21d13 exhibit environmentally dependent regulation and, perhaps as a result, this module is found only under drought stress. In contrast, in modules that have fewer pairwise gene-gene regulatory variations, or when gene expression states are transmitted faithfully from upstream genes in the network, these modules may be more likely observed under all sampled conditions.

Genetic variants affecting expression, whether constitutive differences between varieties (in our study, those with an only significant G effect) or GxE, may be acted on by selection if they ultimately have an effect on phenotype. Past work addressed the possibility of spatial separation across a network between a genetic variant and its effect^35,62^, or between the perception of an environmental stimulus and the downstream transcriptional response^63^. The network context of a genetic variant is important for evolutionary studies because the number and functions of the genes which interact with a variable gene influences the pleiotropy of that genetic variant^64^. Recent work considered the network topology or pleiotropy of gene expression in the context of the evolution of plasticity^48,65^; however, most studies do not account for the prospect of pleiotropic effects of these genetic variants and, as a result, cannot account for variation in the network topology itself. We specifically consider such regulatory interaction variation here. Our large RNASequencing dataset also allowed us to estimate gene co-expression independently in each combination of genotype and environmental treatment, thus relaxing the assumption of an invariant network topology across conditions used previously^48,65,66^. We note that other approaches to account for network structure, such as mapping covariance or principle components as traits in QTL analysis ^3,11,67^ may detect similar regulatory structure.

Changes in gene-gene interactions are found to be clustered in some modules, suggesting a spatial pattern of regulatory variation in networks and supporting the hypothesis that modularity allows for reduced pleiotropic effects of mutations^64^. A related hypothesis is that certain regulatory pathways are under weaker purifying selection, thus leading to accumulation of genetic variation in a subset of modules.

Consistent with this latter hypothesis, we observe that variable interactions are more common in modules not associated with essential biological pathways (e.g., not enriched for GO terms such as translation and ATP synthesis processes). The small overlap between variable modules and the chain structure in Figure 4 is representative of these patterns. Previously, we observed a similar pattern in which individual genes exhibiting expression GxE were localized at the periphery of a gene co-expression network under drought^48^. The pattern we observe here is similar, but now at the level of gene-gene regulatory interactions.

While we have highlighted a number of cases where the interactions among genes vary, we do observe several modules that, while conserved among library types, are enriched for genes that respond to stress. Clique 8-6, for example (Figure 4), is comprised of one module from each of the four library types but is significantly enriched for genes that individually show a significant response to soil drying stress in our study. These gene lists are also enriched for GO terms associated with the light reactions of photosynthesis. Thus, while the gene interaction network of the pathway forms a system whose interactions are constant across various conditions and genotypes, this system is nonetheless environmentally responsive. The mechanism of this response – whether via active regulation or more systems-level metabolic differences -- cannot be determined with this type of data but we highlight this difference from the environmentally dependent gene-gene interactions that we observe in, e.g., Clique 3 (Figure 4).

Our study also revealed a functional connection between cellular and whole-plant traits to specific gene regulatory features. First, modules associated with traits or enriched with GO terms represent hypotheses about the transcriptional control of key drought responses. However, even in modules representing housekeeping pathways, fewer than 50% of genes have annotations; our network-based approach therefore represents a novel means to discover gene function. Second, certain traits, like leaf glucose content, might be controlled by several pathways, as we found glucose to be associated with four modules (i.e., the module overlapping with three modules in glycogen biosynthesis process clique, two drought specific modules, and the module enriched with ABA processes; Table 1). We also show how these associations may change in different environments as some of the glucose-associated modules only exist under drought stress. Collectively, these results suggest that our approach represents an unbiased and systematic way of developing hypotheses about the genes and pathways underlying biological processes.

Collectively, our work highlights how natural selection may modify gene regulation networks to have new functions or environmental responses and may also shed light on how we can modify traits through breeding. First, a gene’s expression can be modified at various steps of its regulation, e.g., at its direct regulator or further upstream. Conversely, modification of a gene’s expression to be responsive to an environmental factor will likely require accounting for gene-gene interactions; our study shows one way to identify such targets.

Second, regulatory variation or environmentally responsive features seem to cluster in specific modules. Variants whose effects are restricted to such modules may have fewer pleiotropic effects and thereby be more likely to be retained in populations. Third, it might be possible to change plasticity to certain stressors without having pleiotropic effects in other environments; modules with many GxE edges but few G edges might be fruitful targets to such avoid trade-offs. On the other hand, targeting modifications to modules with E edges but few GxE edges may prevent rank-changing interactions when breeding for a range of environments^19,20^. While our results suggest how natural or artificial selection may shape, or be shaped by, physiological and regulatory responses to the environment, how these factors directly contribute to drought tolerance and lifetime fitness cannot be determined in the chamber growth experiments like those performed here. Future studies need to build the connection from the current transient gene expression and regulation characterization to fitness in the natural environment.

## METHODS

### Experimental growth conditions and drought treatments

The experiment protocol was adapted from a study by Des Marais et al.^68^ Two accessions of *Brachypodium distachyon* (Bd21, Bd3-1) seeds were placed at 4 °C for seven days to synchronize seedling germination. After that, the seeds were planted in Deepot D16H pots (Stuewe&Sons) filled with Profile porous ceramic rooting media (Profile Products, Buffalo Grove, IL). The dry weight of each pot (DW) and its field capacity for water (FC) were recorded. FC was determined by saturating a pot with water and allowing it to drain under gravity overnight and calculating the weight difference from DW; this was used as the basis for the subsequent controlled soil dry down. Plants were grown in growth chambers with 50% humidity, 500 μmol photons m^-2^ s^-1^ light intensity, and 25°C day/20°C night (12h/12h) and bottom watered with tap water (pH=5-6) every other day throughout the experiment, supplemented with DYNA-GRO GROW Liquid Plant Food 7-9-5 (Dyna-Gro, Richmond, CA, USA) over the first 32 days. On day 33 after sowing, control plants were watered to 85% FC every night while dry-down plants were allowed to decrease their soil water content to 50% FC over 7 days (5% decrease each day). The plants were divided into two groups with Group 1 used to characterize plant responses along the dry-down process, and Group 2 used for gene expression and metabolite content.

On days 0 (85%FC), 2 (75%FC), 4 (65% FC), 5 (60% FC), 6 (55% FC), and 7 (50% FC) nine individuals of each genotype from each treatment (control and drying) were used for gas exchange measurements at 3–5 hours after lights-on, non-photochemical quenching (NPQ) at 5-6 hours after lights-on, and destructive sampling for leaf hydraulic potential, leaf metabolites, and total plant biomass at 6-8 hours after lights-on. Our preliminary analyses determined that gas exchange and leaf water potentials of droughted plants began to deviate from water controls on days 5 and 6 of the dry-down; accordingly, we selected day 6 for more detailed analysis. 48-58 replicates of leaf tissue in each genotype and treatment were measured for gas exchange and photosynthesis at 1.5-5 hours after lights-on on day 6 and harvested for RNA extraction and metabolites assay between 6-7.5 hours after lights-on (equivalent to mid-day). Sampling of individuals was randomized to avoid confounding time-of-day harvest with genotype or treatment.

### Physiological and metabolic measurements

Net CO_2_ assimilation rate (A), and stomatal conductance (g_sw_) of the youngest fully expanded leaf of a selected tiller were determined using a Li-6800 (LI-COR Biosciences, Lincoln, NE). The LiCor chamber was set to a flow speed of 200 μmol s^-1^, humidity at 50%, CO_2_ concentration at 400 μmol mol^-1^, fan speed of 10,000 rpm, and photosynthetic photon flux density (PPFD) at 1000 μmol (m^-2^ s^-1^). Leaves were left for 5 min to stabilize in the chamber before each reading. Photochemical quenching (qP), non-photo-chemical quenching (qN), and Nonphotochemical chlorophyll fluorescence quenching (NPQ), PSII maximum efficiency of light adapted leaves (fv’/fm’) were measured on leaves at noon; Parameters follow Li-6800 protocol of Determining PSII efficiency. Leaf chlorophyll content was estimated by taking the average of two leaves measured with MultispeQ V2.0 prior to lights-on in the growth chamber. The leaf water potential was measured by the Scholander Pressure Chamber Model 670 (PMS Instrument Company, Albany, OR) on the youngest fully expanded leaf of sampled plants.

Above-ground tissue was excised at soil level with a razor blade. Roots were cleaned with tap water to gently remove growth media and dried with tissue paper. Fresh tissue biomass was recorded before samples were dried for 48 h at 80°C and then weighed for dry biomass.

Tissue for RNA and metabolites was flash frozen in liquid nitrogen, stored at −70 °C and freeze-dried overnight prior to grinding with bead-beating. 20mg dry weight is utilized for the following assays. Metabolites were extracted using sequential ethanol extractions following established protocols^69,70^. Protein, starch, and high degree of polymerization (HDP) fructan, total free amino acids, ethanol-soluble carbohydrates (glucose, fructose, sucrose, and low degree of polymerization (LDP) fructan), and free nitrate were quantified following Burnett et al^69^. Metabolite data are expressed here per g dry weight, or as glucose equivalents per dry weight for carbohydrates. Technical and analytical replicates were run for all assays.

### Statistical analyses

For the traits collected during dry-down (Supplementary Data Set 9), most traits were fitted into linear mixed models using lm and lmer ^71^ in R^72^. Glucose, fructose were fitted with generalized linear mixed models with Gamma distribution in glm and glmer ^71^, with family=Gamma(link= “inverse”). Accession (two levels), Treatment (two levels), Harvest day (six levels), and their interactions, were modeled as fixed factors and freeze dry session (four levels) and assay batches (five levels) for metabolites were modeled as random factors. Model selection was first performed on the fixed-effect parts of each model, based on maximum likelihood Akaike information criterion (AIC) using the dredge function in the MuMin package ^73^. Random-effects factors of each model are based on maximum likelihood of models using the anova function. Models are shown in Supplementary Table 2. Throughout the dry-down process, the pairwise significance test between treatments for each accession on each harvesting day (Supplementary Figure 1 and 2) was estimated with a t-test (or Z-test for glm, glmer) adjusted by Tukey using emmeans function ^74^ in R based on the model selected. Outliers were removed before model fitting. The results are visualized using ggplot2 and ggpubr^75,76^.

Similarly, leaf trait data collected on Day 6 (Supplementary Data Set 10) were fit with (generalized) linear mixed models using lm, lmer, glm, or glmer, as appropriate, with fixed factors of Accession, Treatment, and their interaction, and random factors of growth batch, freeze dry sessions, and assay batch for metabolites (Figure 1). The p-values shown are based on likelihood ratio test (Chisq test) of dropping each term using anova function.

### Transcriptome sampling and processing

Leaf tissue flash frozen on liquid nitrogen was ground using an MP FastPrep-24™ Classic bead beating grinder with CoolPrep™ 24 x 2mL Adapter (MP Biomedicals) loaded with dry ice. Approximately 100mg of tissue was used for RNA extraction using Spectrum Plant Total RNA Kit (Sigma-Aldrich, USA) according to the manufacturer instructions, followed by DNase treatment (On-Column DNase I Digestion Set, Sigma-Aldrich, USA). The quality and quantity of the RNA samples were determined both by absorbance measurements (Nanodrop 1000, NanoDrop Technologies) and with a Tape Station (4200 TapeStation System, Agilent). 25ng of RNA were processed for High-Throughput 3’ Digital Gene Expression ^77,78^ by the MIT BioMicro Center, pooled and then sequenced across three flowcells on an Illumina Novaseq (NovaSeq6000 Sequencing System, Illumina) generating, on average, 3.9±1.3 million raw reads per sample.

Raw sequence reads were trimmed by Trimmomatic ^79^ and Cutadapt ^80^ to remove adapters and poly(A) tails. All reads were mapped to the *B. distachyon* Bd21 primary transcriptome sequence (Bdistachyon_556_v3.2.transcript_primaryTranscriptOnly, downloaded from Phytozome) and the annotated sequence of the whole Bd21 plastome, which is available at Genbank (LT558597 accession). Read alignments from Bowtie2 ^81^ were corrected by RNA-Seq by Expectation-Maximization (RSEM) ^82^ to account for non-uniquely aligned reads. Bowtie2 alignment is done with seedlength -L 25 and mismatch allowance -n 2 which allows mismatches of Bd3-1 read alignments. Alignment with -- estimate-rspd specifies the reads position distributions that are highly 5’ or 3’-biased ^82^. Reads with the same unique molecular identifier and aligned to the same genes were removed to reduce PCR biases during library generation ^77,78^ prior to adjustment in RSEM. RNASeq reads from Bd21 and Bd3-1 mapped to the Bd21 references at 77.0±1.7% and 78.1±1.8 %, respectively, suggesting no genotypic effect on mapping.

Reads which did not map to the genome during this first round were mapped onto ribosomal RNA, after which the total mapping rates increased to ∼96%.

### Differential gene expression (DGE)

RNASeq reads were normalized using Relative Log Expression normalization implemented by DESeq2 ^83^; these normalized data with log_2_ (Count+1) transformation are used for downstream analyses. The grouping is visualized with PCA using factoextra package in R.^84^ In DGE step, genes with >0 counts in more than 10 samples are kept. DESeq2 fit the transcript count of each gene into the generalized linear model Transcript Count ∼ Genotype + Treatment (+ Genotype * Treatment) or Transcript Count ∼ Treatment, stratified by genotype or Transcript Count ∼ genotype, stratified by treatment. Each gene is fitted into a negative binomial distribution. Results in Figure 3 were visualized with ggplot in R. ^75,76^

### Identifying patterns of variation in clusters of gene-gene interactions

For this and following analyses, we removed transcripts exhibiting low variance (<0.1) among replicates in each library type; in total, 16,207 genes are included. Co-expression modules in each library type (Bd21 Control, Bd21 Drought, Bd3-1 Control, Bd3-1 Drought) were generated using Weighted Gene Co-expression Network Analysis (WGCNA)^57^. For WGCNA, the starting correlation matrix is unsigned, as both negative and positive correlations are of interest in this study. Weight power=5 is added to the correlation matrix to achieve a scale-free adjacency matrix. The topology similarity matrices in the Bd21c, Bd3-1d, and Bd3-1c data sets are scaled to have a similar distribution to Bd21d, which we chose arbitrarily as the reference library type.

Dendrograms were generated from topology similarity matrices, and modules were generated by cutting at two parameter settings: (1) moderate size modules, used for most analysis below, high dissimilarity (height= NULL(0.99) with a dynamic hybrid algorithm (minClusterSize =30, deepsplit=2, pamStage = FALSE, mergeheight=0.25); (2) conservative (smaller) modules, used for causal inference analyses discussed below, cutHeight = 0.984, minClusterSize = 15, deepSplit = 3, pamStage = FALSE, mergeheight=0.25. The two different settings are used to better understand the effect of threshold choice within each library type. In brief, our WGCN analyses recovered 15 and 24 modules in Bd21c in the two settings, respectively; 14 and 18 modules in Bd21d in the two settings, respectively; 14 and 16 modules in Bd3-1c in two settings, respectively; and 15 and 20 modules in Bd3-1d in the two settings, respectively. Gene membership is summarized in Supplementary Data Set 9.

We next addressed metrics of module preservation across our four library types, Bd21c, Bd21d, Bd3-1c, and Bd3-1d. First, we assessed the degree to which the lists of genes in a given module are conserved among our two genotypes and two environmental conditions using a test for membership overlap. We required that module overlap must be significant in all two parameter settings (moderate and conservative, described above, as shown in Figure 4, three modules with unstable overlapping are not shown). To examine the overlap in gene module membership among library types we used the SuperExact test ^58^ and a p-value cut-off of with < 10^-12^ with the alternative hypothesis that the size intersect between two groups or among more groups is larger than expected by chance (Supplementary Data Set 5). The cut off is chosen to ensure a p-value <0.05 after applying Bonferroni correction of multiple tests. Hypergraphs representing module overlap networks -- here, termed “cliques” for subnetworks -- were visualized using the R package igraph ^85,86^.

Second, we tested for enrichment of transcripts exhibiting significant DGE in our co-expression modules using Fisher’s Exact Tests ^58^. Significant enrichments (p-value <0.001 uncorrected by test number) are reported in Supplementary Data Set 6 and summarized in Table 1. Here the enrichment is reported for modules estimated from our conservative parameter settings to be consistent with following network inference analysis.

Third, biological roles of each module were inferred using GO term enrichment, by inspecting gene annotations, and by testing for module correlation with the sampled physiological and metabolic traits. Module GO enrichment is conducted on using the PANTHER classification^87^ and implemented in rbioapi^88^, with Fisher’s exact statistical overrepresentation tests of GO biological process. Only the significantly enriched GO terms at FDR = 0.1 and the lowest-rank GO categories are reported. Here the enrichment is reported if positive in modules of any of the parameter settings (Supplementary Data Set 7). For common GO term comparison among modules in an overlapping clique, all GO categories are considered (summarized in Table 1). Common genes and annotations in overlapping cliques are list in Supplementary Data Set 3-4.

Gene annotations are from MapMan (https://mapman.gabipd.org/mapmanstore) based on the Phytozome database. Correlations between module eigengenes (i.e., their first principal component) and traits was performed in each library type. The correlation p-values are reported in Supplementary Data Set 11 and summarized in Table 1 based on moderate criteria modules, uncorrected for the number of tests.

Finally, we tested the hypothesis that genes within modules share common DNA motifs corresponding to known transcription factor binding sites. All possible 6-mer sequences were tested for enrichment in the −500 to +200 upstream window regions^89^ relative to each gene’s predicted transcriptional start site in the Bd21 reference genome, in the module using AME in MEME suite^90^. The AME parameters were: --verbose 1 -- scoring avg --method fisher --hit-lo-fraction 0.25 --evalue-report-threshold 10.0, with the control set based on all the genes. The Arabidopsis DAPSeq database ^91^ was used as a reference to interpret enriched motif sequence identified. Here the enrichment is reported for moderate criteria modules.

### Causal network inference in modules

Methods such as WGCNA, based on correlations in transcript abundance, cannot provide information about the direction of regulatory interactions. Our high replication provides the power to model genes as random variables and infer conditional independence of gene expression to represent possible causality. Here, we use the Unknown-Targeted Interventional Greedy Sparest Permutation (UT-IGSP) algorithm ^92^ to infer regulatory causality, representing a group of genes as a directed acyclic graph (DAG). We use all genes in a given WGCNA module – estimated from one library type -- as priors to reduce latent variables in the model, relying on the simplifying assumption that there are no edges among genes between modules. We include both samples that the module is inferred from (here, representing “observational” data) as well as samples from all other library types (here, representing “interventional data”), representing the assumption that most of the gene regulation structure is the same between library types. In these analyses, we use the conservative module criteria (most restrictive inclusion in the WGCNA analyses) and limit the analysis to modules with number of genes fewer than 70% of samples in the specific library type. When testing for modules that are conserved within a treatment or within a genotype, modules with as many as 70 genes can be used because more samples are available for inference. We use stability selection ^93^ as implemented in Belyaeva et al ^94^ to minimize the effect of hyperparameters. Edges with a selection probability higher than upper 0.95 (or 0.97 for bigger conserved modules to keep sparse edges, and 0.95 for modules with glucose as a node) quantile are kept. Genes not found to be connected to other genes are not shown. DAGs were exported from R using the rgraphviz and visualized using Graphviz Visual Editor ^95^.

### Identifying patterns of variation in gene-gene interactions

We extended our analysis of gene expression responses using a regression approach that explicitly models the magnitude and direction of putative interaction between genes based on their transcript abundances. For a putative regulator and target pair identified in our DAGs, we regress the target gene transcript abundance on the regulator transcript abundance, and test whether the slope and intercept coefficients of regression changes as a function of genotype or environmental treatment. When there are multiple inferred regulators for a target gene, a multivariate linear regression is applied.

For the hypothesis that gene *x_i_* (*i* = 1, …, *n*)) regulates gene *y*, to infer the intercept or slope differences between genotypes, treatment, or their interactions, we modeled their (log_2_-transformed) gene expression *x_i_* and *y* as:

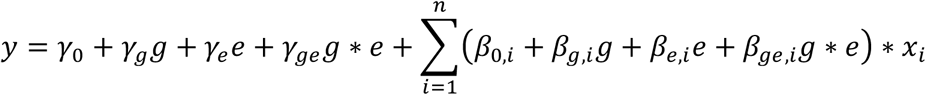

Here *g,e* are used to differentiate the dataset used to train the parameters (i.e., *g, e* are “0,0”, “0,1”, “1,0”, “1,1” for Bd21 control, Bd21 drought, Bd3-1 control and Bd3-1 drought respectively). Thus, consistent with the DGE test using DESeq2, above, a significant p-value of γ_g_, β_g_ suggest a genotypic effect on regression intercept and slopes, γ_e_, β_e_ suggest an environmental effect on intercept and slopes, and γ_ge_, β_ge_ suggest a GxE effect of intercept and slopes. Only genes with regression R^2^ greater than >0.5 are considered, as estimated with the Car package in R. ^96^ P-value of <0.05 is reported to be significant without multiple test correction. Note that if the transcript abundances are standardized before fitting our regression models, the slope differentiation test should closely match the results of CILP tests introduce by Lea et al^67^ which we include as Supplementary Figure 6.

### Allele specific expression

We used allele-specific expression to assess whether GxE in transcript abundances were caused by *cis-* or *trans-*acting variants. Bd21 served as a pollen donor on emasculated Bd3-1 flowers, resulting in ten F1 hybrid seeds. To confirm successful outcrossing in this largely selfing species, we extracted DNA from leaves (DNeasy Plant Pro Kit; Qiagen, MD, US) and tested for a known length polymorphism on a 3% agarose gel. We performed a dry-down experiment on these F1s and extracted leaf RNA, as above. 100ng of RNA for each of 10 samples along with 12 samples (three replicated of each genotype and condition) from the initial experiment were aliquoted for sequencing. For DGE, and to identify SNPs between alleles, we generated libraries using NEB Ultra II Directional RNA with Poly(A) Selection (New England Biolabs, Ipswich, MA). Libraries were sequenced on an Illumina Novaseq generating 34.1 ± 13.7 million raw reads per sample.

To generate a reference diploid transcriptome, RNAseq reads of each Bd3-1 sample were aligned to the Bd21 reference genome (Bdistachyon_556_v3.2) based on a RNAseq short variant discovery pipeline ^97^, with STAR alignment ^98^ of two pass and alignSJoverhangMin=30 and alignSJDBoverhangMin=10, variants calling GATK ^99^ and the flag of dont-use-soft-clipped-bases. The variants called in each sample were then filtered to find SNPs and in/dels with homozygous alternative genotype mapping. To reduce false discovery only variants identified in more than half of the replicates of each genotype were kept. The synthesized Bd21 or Bd3-1 vcf together with Bd21 genome and annotation gff was then used to generate *de novo* Bd21 or Bd3-1 genome using g2gtools ^100^, and a diploid genome with emase ^101^. The reads from all the samples were mapped to this diploid transcriptome using bowtie^102^ and a count matrix was estimated from alignments using emase (an alternate pipeline employing bowtie2 followed by RSEM provides similar results; not shown) to account for the multi-mapping particularly accounting for allelic expression. Isoforms are not differentiated in this study.

Genes with different counts between two alleles are considered to have sequence variants but sparse differences in a gene might not accurately differentiate two alleles. Accordingly, we included 3963 genes that have more than >1 count of the allele in one accession and <20% total count of the allele in the opposing accession in all samples, modified from Gould et al ^103^. The RNASeq count matrix was normalized using RLE in DESeq2, as above. All genes (>0 in more than 10 samples, including allele-indifferentiable genes) are included in the normalization step with allele-distinguishable genes shown as allelic expression and other genes as total expression to ensure the normalization factors are calculated based on most genes for the comparable of genes with allelic variants. Possible read-mapping bias was assessed as in Lovell et al. ^104^ shown in Supplementary Figure 7. PCA of samples are shown in Supplementary Figure 8.

We employed a pipeline based on DESeq2 as introduced by Lovell et al ^104^. Briefly, the transcript abundance of a target gene is fit into full model

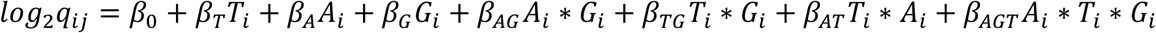

where *q_ij_* is gene expression of sample i for gene j, *T_i_* is treatment (control: 0, drought: 1), *A_i_* is allele type (Bd21: 0, Bd3-1: 1), *G_i_* is generation (F1: 0, Parents: 1). Accordingly, β_A_ tests for *cis-*acting regulation; β_AG_ tests for *trans-*acting; β_AT_ tests for *cis-*by-environment-acting; and β_AGT_ tests for *trans-*by-environment-acting. Here, the approach was also modified to perform tests in each environmental condition (as a “half model”). To be specific, in half model, each test of cis/trans is done separately in each condition because of our limited power and possible low expression of the gene in samples of some conditions; cis/trans by environment interaction cannot be directly tested in this analysis. P-values <0.1 is reported to be significant without multiple test correction. Results are summarized in Supplementary Data Set 12.

## Supporting information

Supplementary Data Sets 1-12

## DATA AVAILABILITY

The datasets and computer code produced in this study are available in the following databases:

- RNA-Seq data: Gene Expression Omnibusm GSE267880 (https://www.ncbi.nlm.nih.gov/geo/query/acc.cgi?acc=GSE267880)
- RNA-Seq data for allele specific expression: Gene Expression Omnibusm GSE267878 (https://www.ncbi.nlm.nih.gov/geo/query/acc.cgi?acc=GSE267878)
- Analysis scripts: GitHub (https://github.com/jyjiey/GxE_in_gene_regulation_network).

## SUPPLEMENTARY DATA

**Supplementary Data Set 1.** Gene DGE list

**Supplementary Data Set 2.** GxE DGE gene GO enrichment

**Supplementary Data Set 3.** Module common genes

**Supplementary Data Set 4.** Module common gene annotation sorted

**Supplementary Data Set 5.** Module overlap p-value and intersect gene number

**Supplementary Data Set 6.** Module G,E,GxE DGE gene enrichment p-value

**Supplementary Data Set 7.** Module GO enrichment

**Supplementary Data Set 8.** Module traits correlation p-value

**Supplementary Data Set 9.** Traits data along dry-down

**Supplementary Data Set 10.** Traits data on day6 matching RNA samples

**Supplementary Data Set 11.** Gene module membership

**Supplementary Data Set 12.** ASE type

## ACKNOWLEDGEMENTS

This work was supported by a grant from the MIT Jameel Water and Food Systems Lab (JWAFS) and NSF CAREER Award (IOS 2239070) to DLD. JY was supported, in part, by a graduate research fellowship from MIT JWAFS. ACW & AR were supported through the United States Department of Energy contract no. DE-SC0012704 to Brookhaven National Laboratory. We acknowledge the MIT SuperCloud and Lincoln Laboratory Supercomputing Center for providing (HPC, database, consultation) resources that have contributed to the research results reported within this paper. We also acknowledge the computational support of Massachusetts Green High-Performance Computer the Commonwealth Computational Cloud for Data Driven Biology (C3DDB).

## AUTHOR CONTRIBUTIONS

DLD and JY designed the research; JY, ACB, AR and DLD performed research; JY and DLD design the analytic and computational tools, analyzed data. DLD and JY wrote the paper with contributions from AR and ACB.

## SUPPLEMENTARY INFORMATION

**Supplementary Figure 1.**
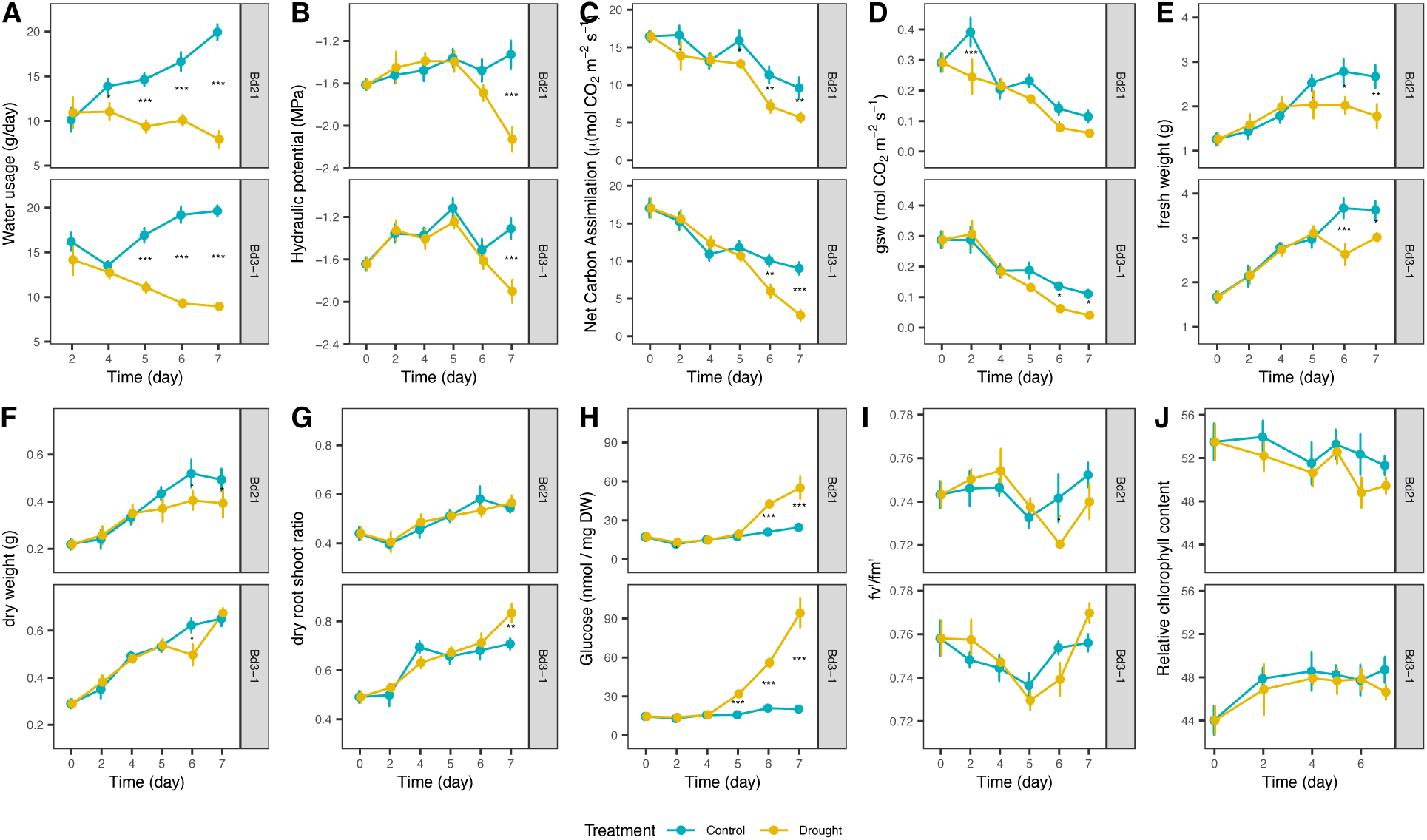
Comparison of physiological and metabolic traits between control and dry-down in two accessions from Day 0 to Day 7. (Generalized) Linear models were determined based on AIC score. Random-effects components of the models were based on likelihood ratio tests. Differences of means shown here with asterisks, were determined using contrasts between the two treatments at each time point in each genotype by Tukey’s test. N = 9 for each point.

**Supplementary Figure 2.**
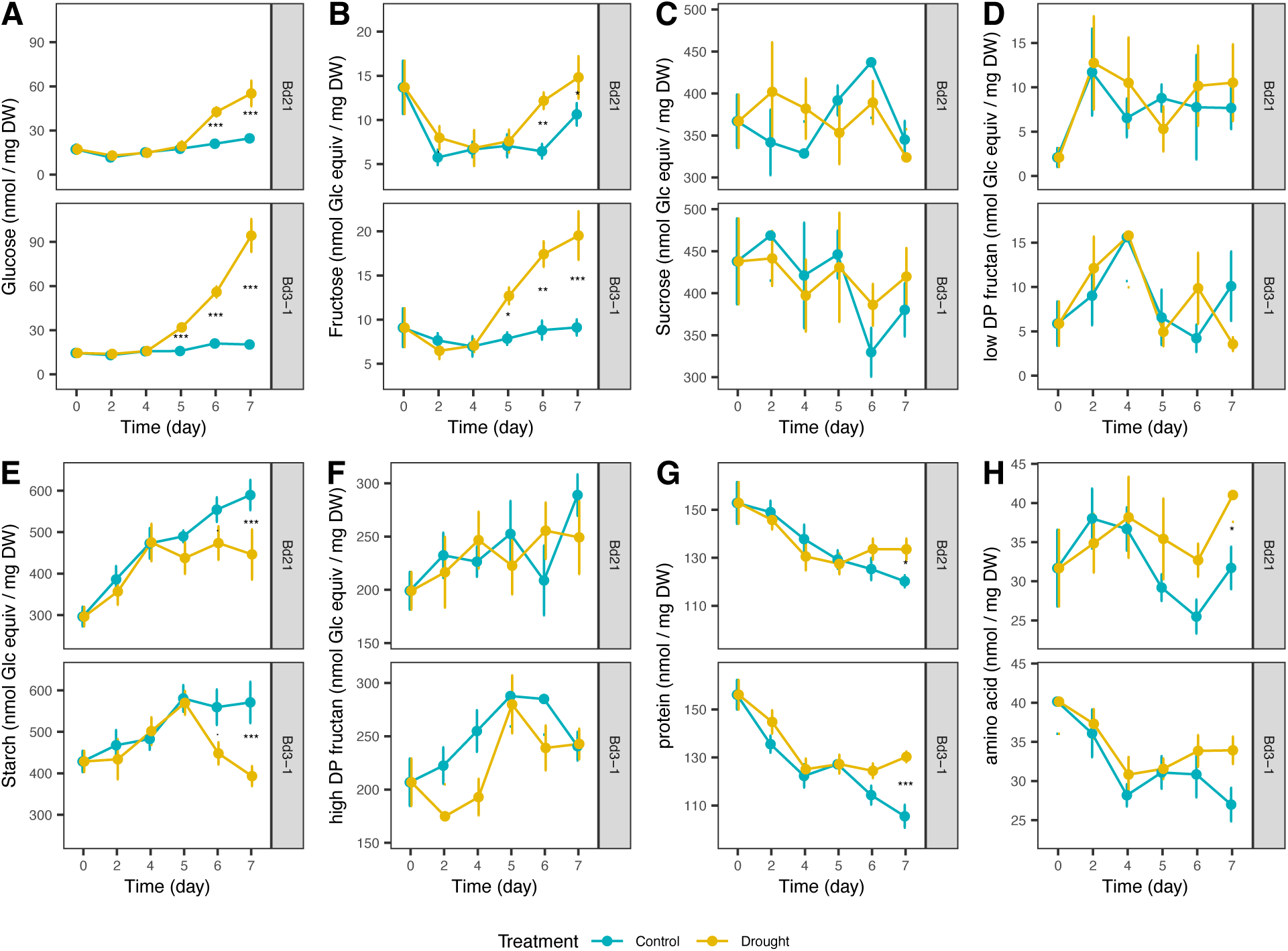
Other physiological traits during dry-down in addition to Supplementary Figure 1. Comparison of physiological and metabolic traits between control and dry-down in two accessions from Day 0 to Day 7. (Generalized) Linear models were determined based on AIC score. Random-effects components of the models were based on likelihood ratio tests. Differences of means shown here with asterisks, were determined using contrasts between the two treatments at each time point in each genotype by Tukey’s test. N = 9 for each point.

**Supplementary Figure 3.**
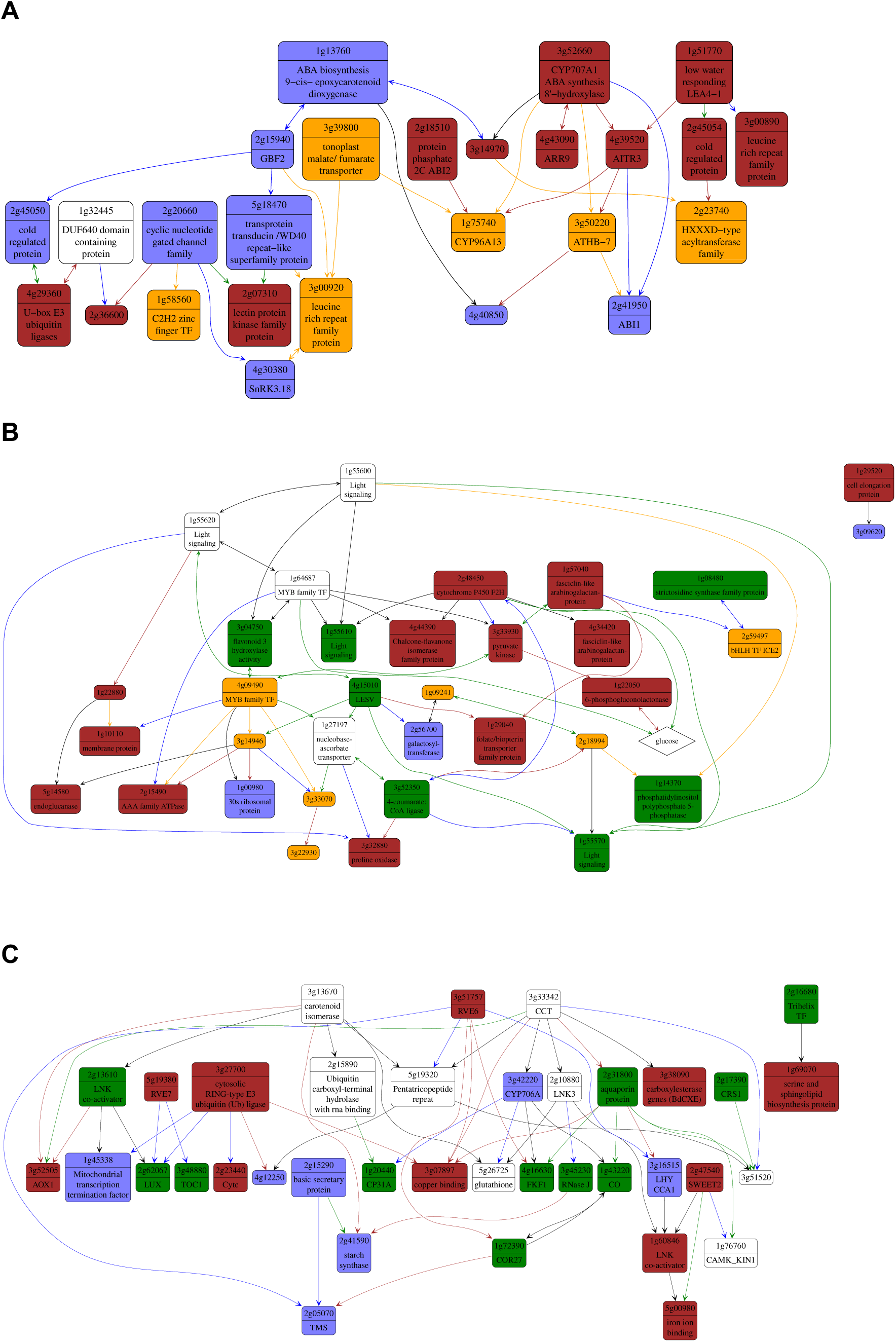
Inferred directed interactions within selected modules with gene DGE and edge DGE changes, corresponding to **A)** Module 12 in Bd21d, **B)** Module 14 in Bd3-1d and **C)** Module 10 in Bd21d shown in Figure 5, with node colors are based on gene DGE and edge colors are based on edge DGE changes (blue: E, green: G, brown: G+E, orange: GxE, black/white: none).

**Supplementary Figure 4.**
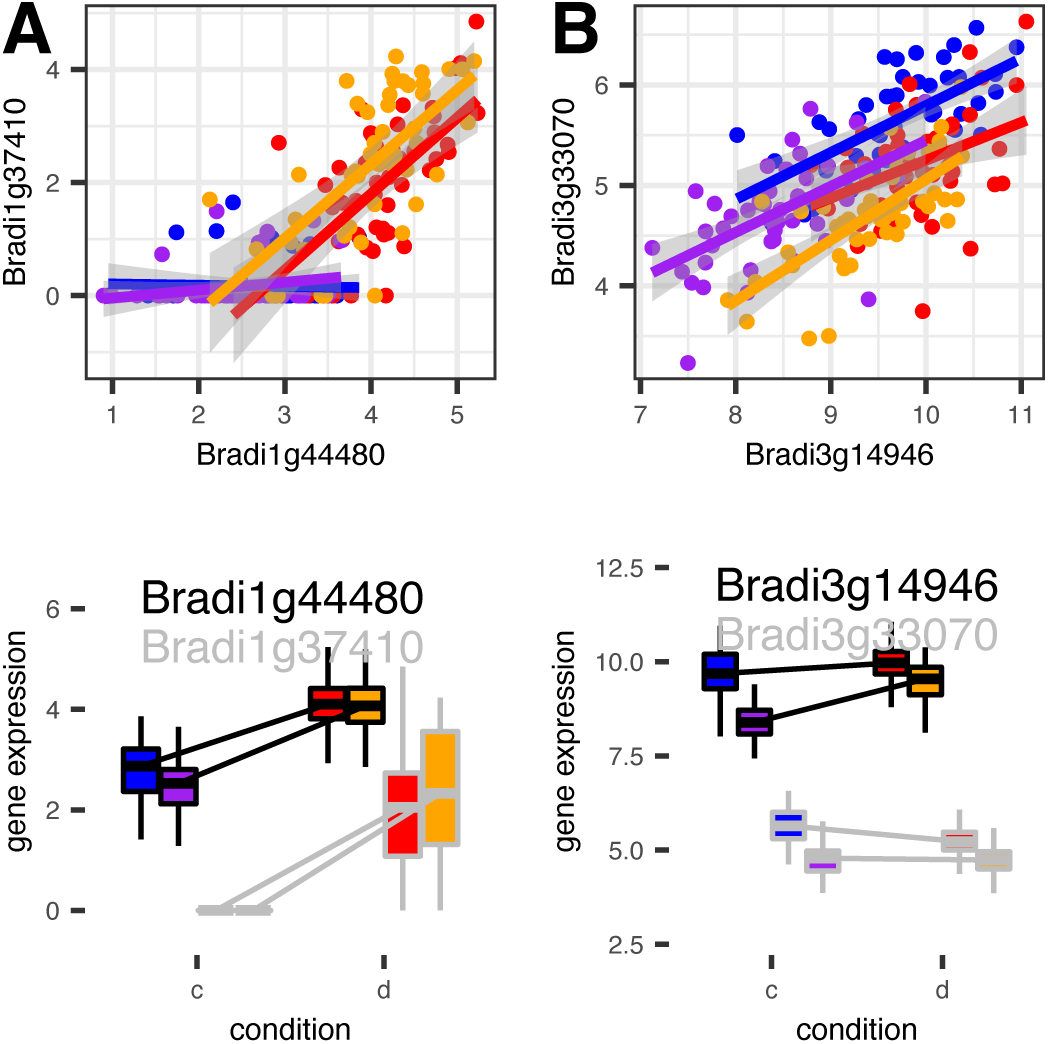
Examples of regulations between pairs of genes selected in addition to Fig.6 A) shows regulation E slope changes, and B) shows transmission of GxE DGE. They are illustrated with linear regression between regulator and target, their expression levels based on log2(norm_count+1). Bd21c, Bd21d, Bd3-1c, Bd3-1d are colored with blue, red, purple, orange, respectively.

**Supplementary Figure 5.**
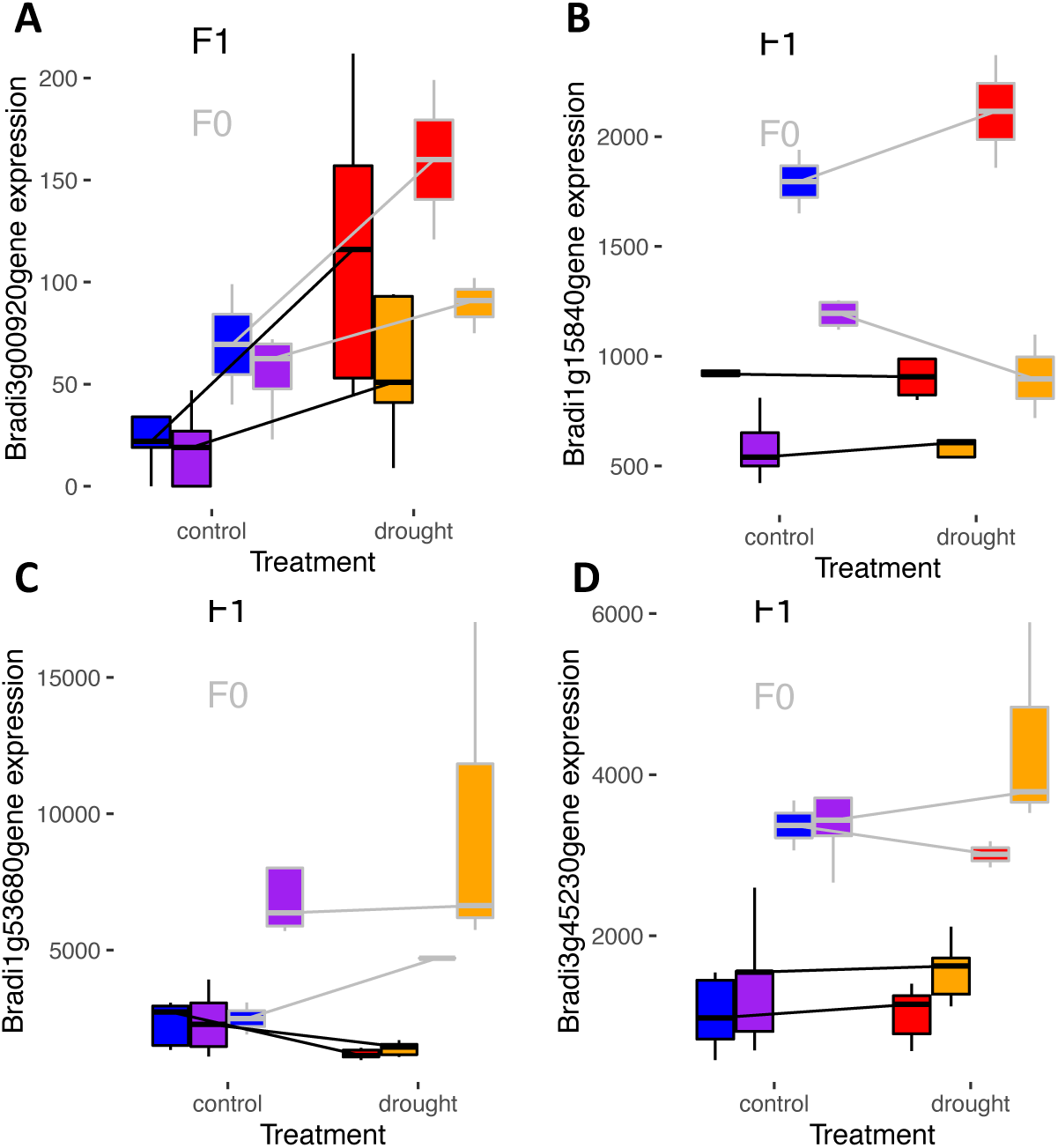
ASE of the genes in Figure 6 (Bradi3g00920, Bradi1g15840, Bradi1g53680, Bradi3g45230). F1 is expression in hybrid and F0 is expression in parents. Values are normalized count. Bd21c, Bd21d, Bd3-1c, Bd3-1d are colored with blue, red, purple, orange, respectively.

**Supplementary Figure 6.**
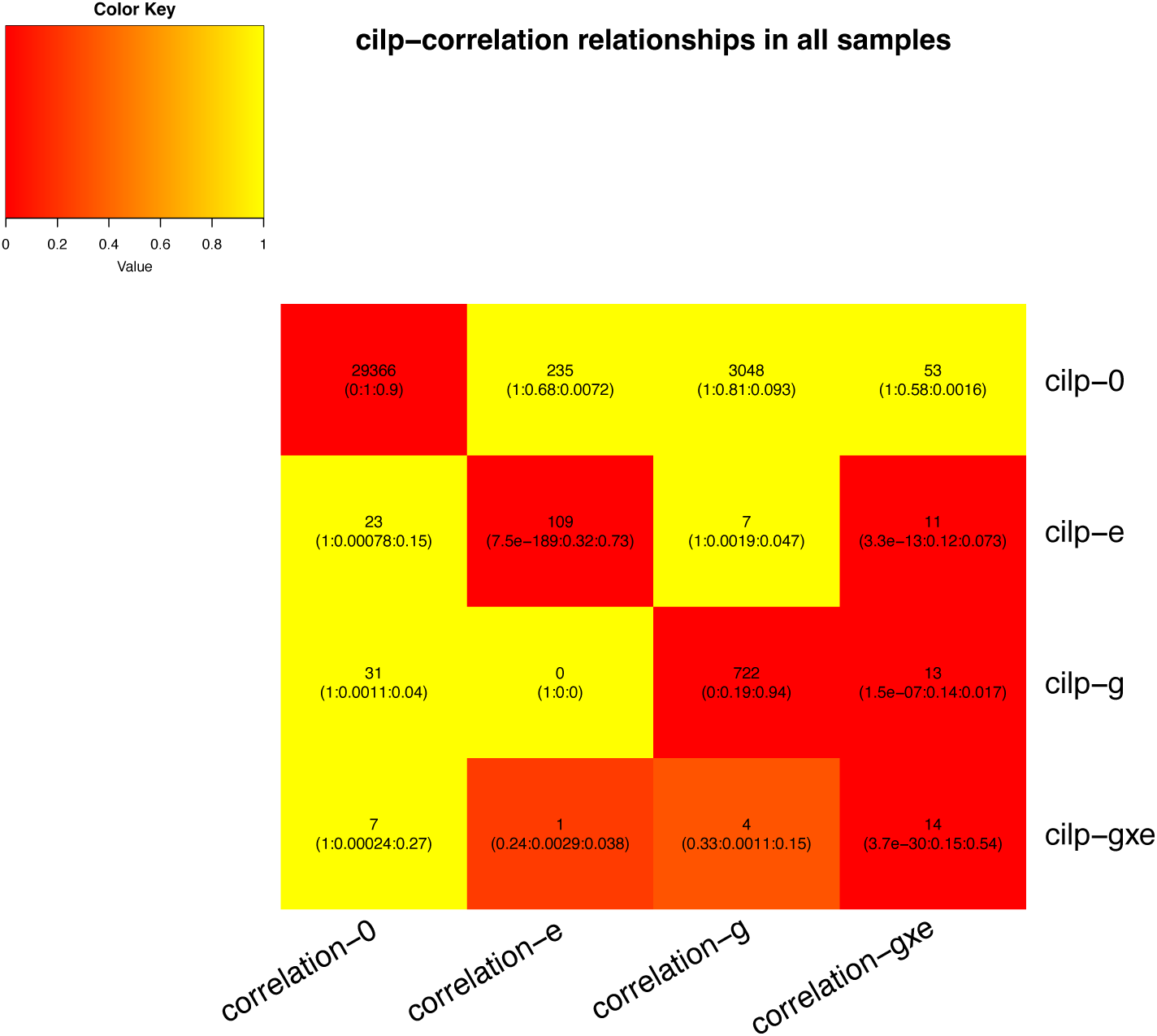
Heatmap comparing CILP and correlation variation from regression method based on gene pairs with correlation >0.75 in any of the library types. The number represent the # of the cases found in the intersect. The three number in the parenthesis are p-value of fisher’s exact test, the proportion of the genes in the intersect in the correlation category, and the proportion of the genes in the intersect in the CILP category. The color is based on p-value.

**Supplementary Figure 7.**
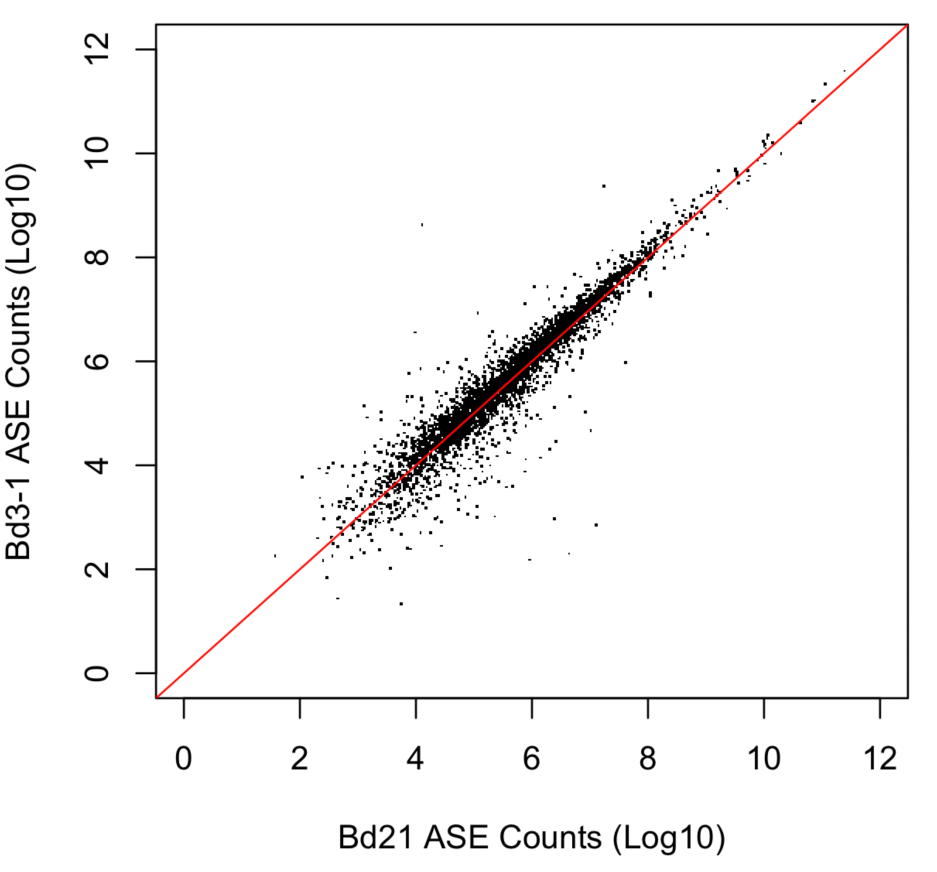
mean reads of Bd21 allelic count vs. mean reads of Bd3-1 allelic count across all samples of allele specific genes suggest no alignment bias to Bd21 or Bd3-1.

**Supplementary Figure 8.**
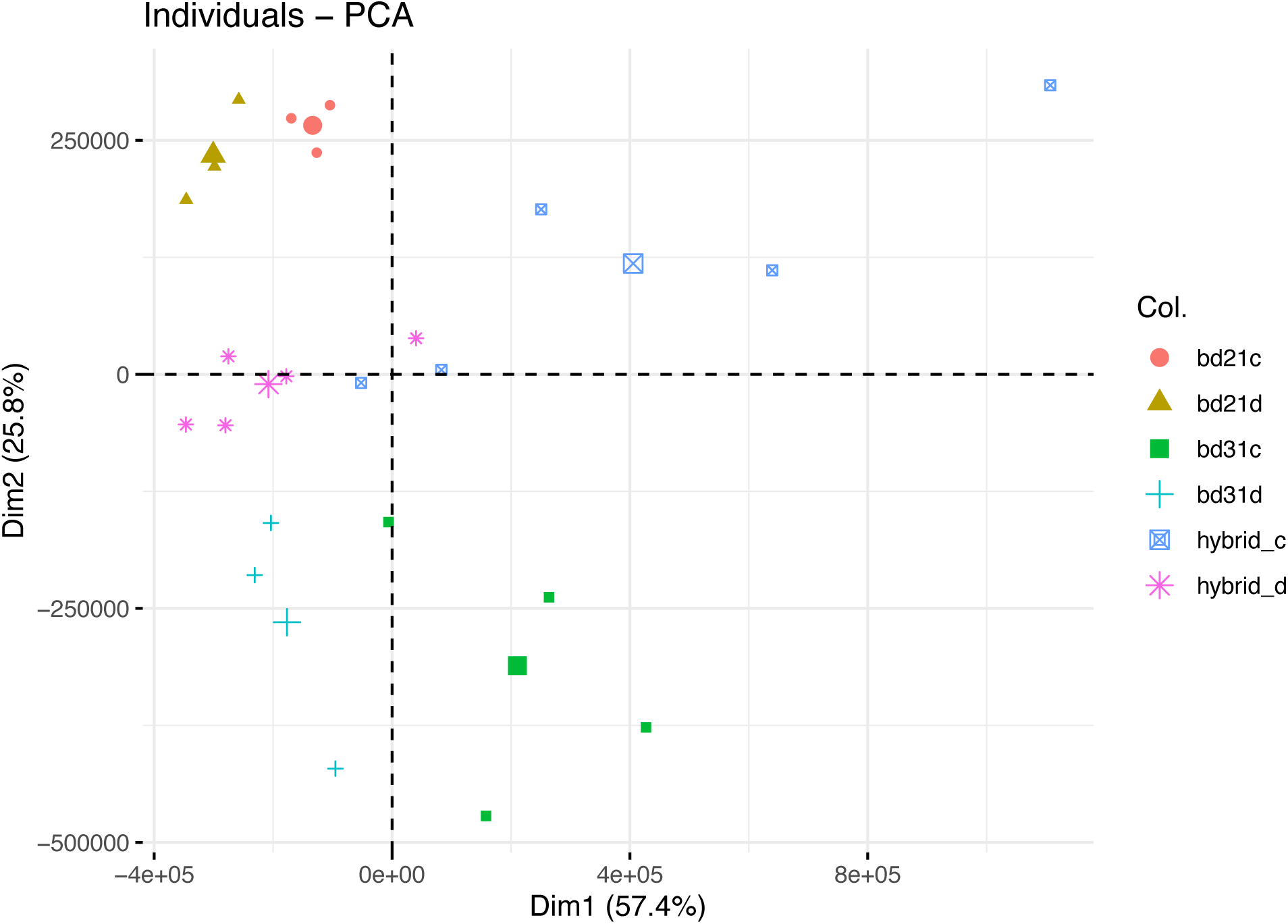
PCA of allelic gene counts colored based on samples genotype and treatment (after normalization)

**Supplementary Table 1.** Percentage of E, G, G+E and GxE DGE change, and slope change identified in edges or intercept change in genes in small module (<40 genes) inferred gene regulation network (e.g., Figure 4).

| module no. | total edges | edge DGE change |  |  |  | edge rewiring % |  |  |  | total genes | gene intercept change % |  |  |  |  |
| --- | --- | --- | --- | --- | --- | --- | --- | --- | --- | --- | --- | --- | --- | --- | --- |
|  |  | G | E | G+E | GxE | G | E | G+E | GxE |  | E | G | GxE | G+E | Non |
| bd21c_13 | 67 | 25 | 4 | 10 | 6 | 10% | 12% | 3% | 9% | 33 | 3% | 24% | 24% | 6% | 39% |
| bd21c_14 | 52 | 19 | 8 | 5 | 0 | 8% | 17% | 0% | 6% | 32 | 6% | 25% | 13% | 9% | 28% |
| bd21c_17 | 37 | 16 | 4 | 6 | 0 | 5% | 5% | 3% | 5% | 27 | 11% | 11% | 15% | 19% | 33% |
| bd21c_21 | 30 | 8 | 4 | 12 | 0 | 10% | 20% | 3% | 7% | 19 | 5% | 5% | 21% | 21% | 37% |
| bd21c_22<br>(part of 9) | 22 | 5 | 3 | 3 | 0 | 5% | 23% | 9% | 5% | 18 | 6% | 22% | 17% | 0% | 28% |
| bd21c_23<br>(part of 10) | 25 | 5 | 3 | 10 | 0 | 4% | 8% | 4% | 0% | 18 | 6% | 6% | 17% | 22% | 33% |
| bd21d_12 | 41 | 5 | 11 | 12 | 11 | 34% | 15% | 5% | 12% | 29 | 3% | 17% | 17% | 7% | 31% |
| bd21d_13 | 32 | 2 | 11 | 7 | 5 | 44% | 3% | 6% | 6% | 25 | 0% | 4% | 32% | 8% | 16% |
| bd21d_16<br>(part of 5) | 23 | 6 | 9 | 6 | 1 | 17% | 4% | 4% | 17% | 20 | 5% | 15% | 5% | 25% | 30% |
| bd31d_12 | 68 | 15 | 16 | 11 | 6 | 10% | 9% | 0% | 1% | 37 | 8% | 8% | 27% | 24% | 11% |
| bd31d_14 | 51 | 17 | 3 | 8 | 0 | 10% | 4% | 4% | 4% | 28 | 0% | 25% | 14% | 21% | 25% |
| bd31d_18<br>(part of 21) | 32 | 4 | 10 | 10 | 2 | 13% | 19% | 9% | 9% | 19 | 11% | 5% | 16% | 21% | 37% |
| bd31d_19 | 108 | 44 | 15 | 18 | 4 | 7% | 24% | 5% | 6% | 33 | 6% | 12% | 15% | 21% | 39% |
| bd31c_13 | 79 | 8 | 4 | 0 | 0 | 8% | 5% | 3% | 13% | 31 | 19% | 10% | 10% | 10% | 48% |
| bd21c_5<br>(intersect with<br>Bd21d_2) | 65 | 25 | 9 | 8 | 0 | 5% | 18% | 2% | 9% | 36 | 14% | 8% | 8% | 17% | 39% |
| bd31c_10 | 55 | 11 | 19 | 10 | 0 | 7% | 9% | 2% | 11% | 33 | 12% | 15% | 21% | 27% | 21% |
| bd31c_16 | 27 | 13 | 1 | 3 | 0 | 15% | 19% | 0% | 15% | 15 | 7% | 20% | 7% | 27% | 33% |

**Supplementary Table 2.**
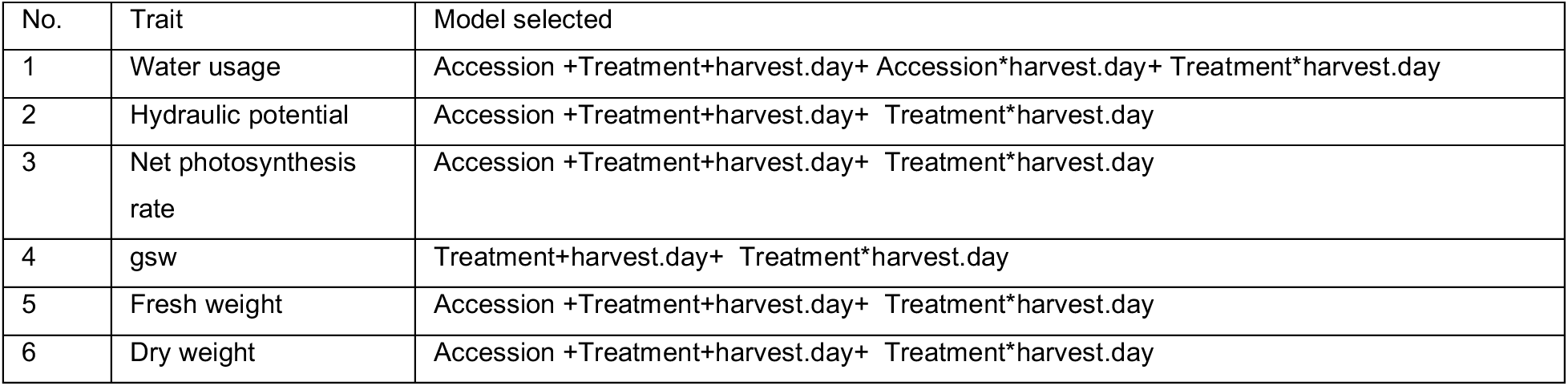
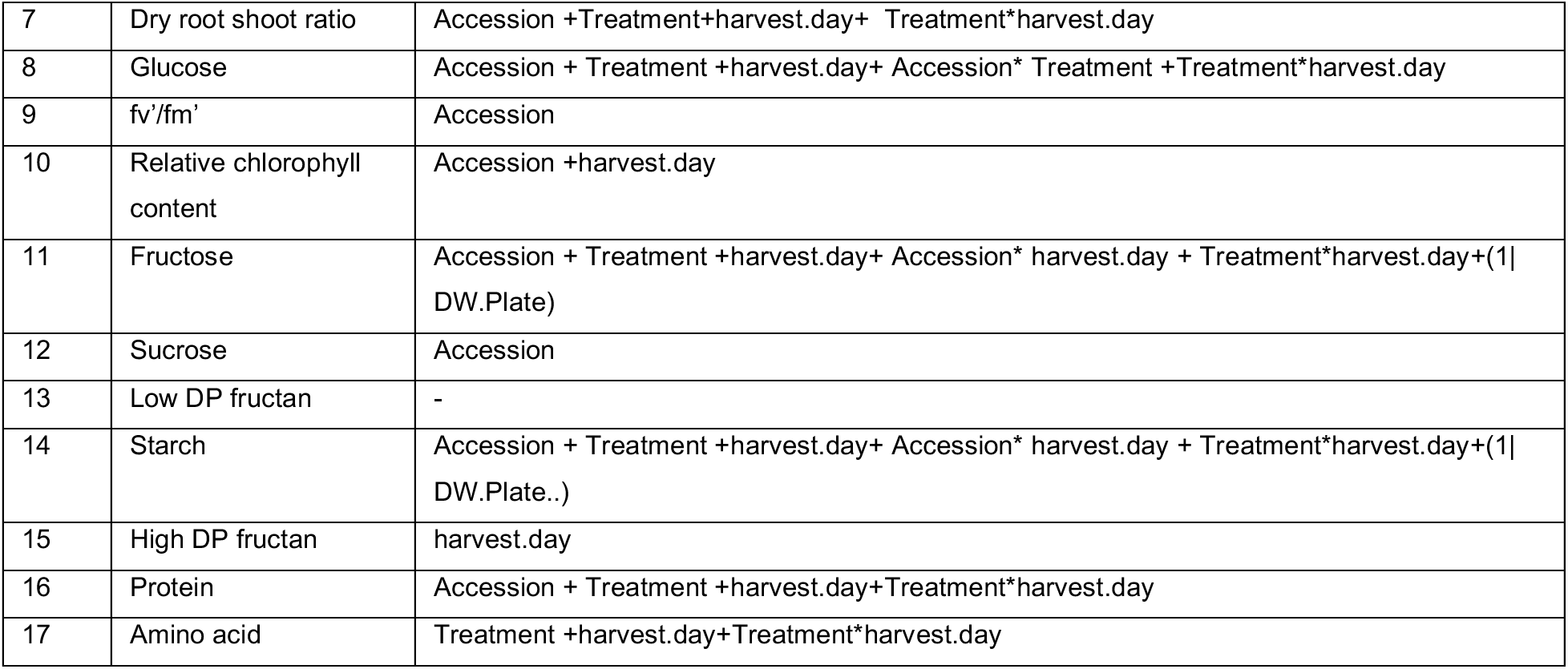
Models used for physiological traits during dry-down reported in both Supplementary Figure 1 and 2. (Generalized) Linear models were determined based on AIC score. Random-effects components of the models were based on likelihood ratio tests.

### Supplementary File 1 Direct test of DGE change between two genes

#### Direct test of DGE change between two genes

We fit pairs of genes in regressions to formally test whether DGE changes occur between them. To be specific, a vector of log2-transformed transcript abundance for two correlated genes were fit with the following model:

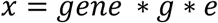

where *x* is a combined vector of gene expression of two genes, and variable *gene* (1,2) differentiates which the gene of gene expression value *x*. Thus, the regression coefficients of *gene* g* which differ significantly from 0 suggest that effects of genotype on transcript abundances differ between the two genes, and similarly coefficients of *gene* e* which differ significantly from 0 suggest that effects of treatment environment on transcript abundances differ between the two genes, and significant coefficients for *gene* g * e* suggests a GxE effect on transcript abundance. P-values <0.001 is reported to be significant without multiple test corrections. Results are shown in Supplementary Figure 2. This method avoids the indirect comparison by DGEs of two genes, but generally consistent with it (see Supplementary Figure 3).

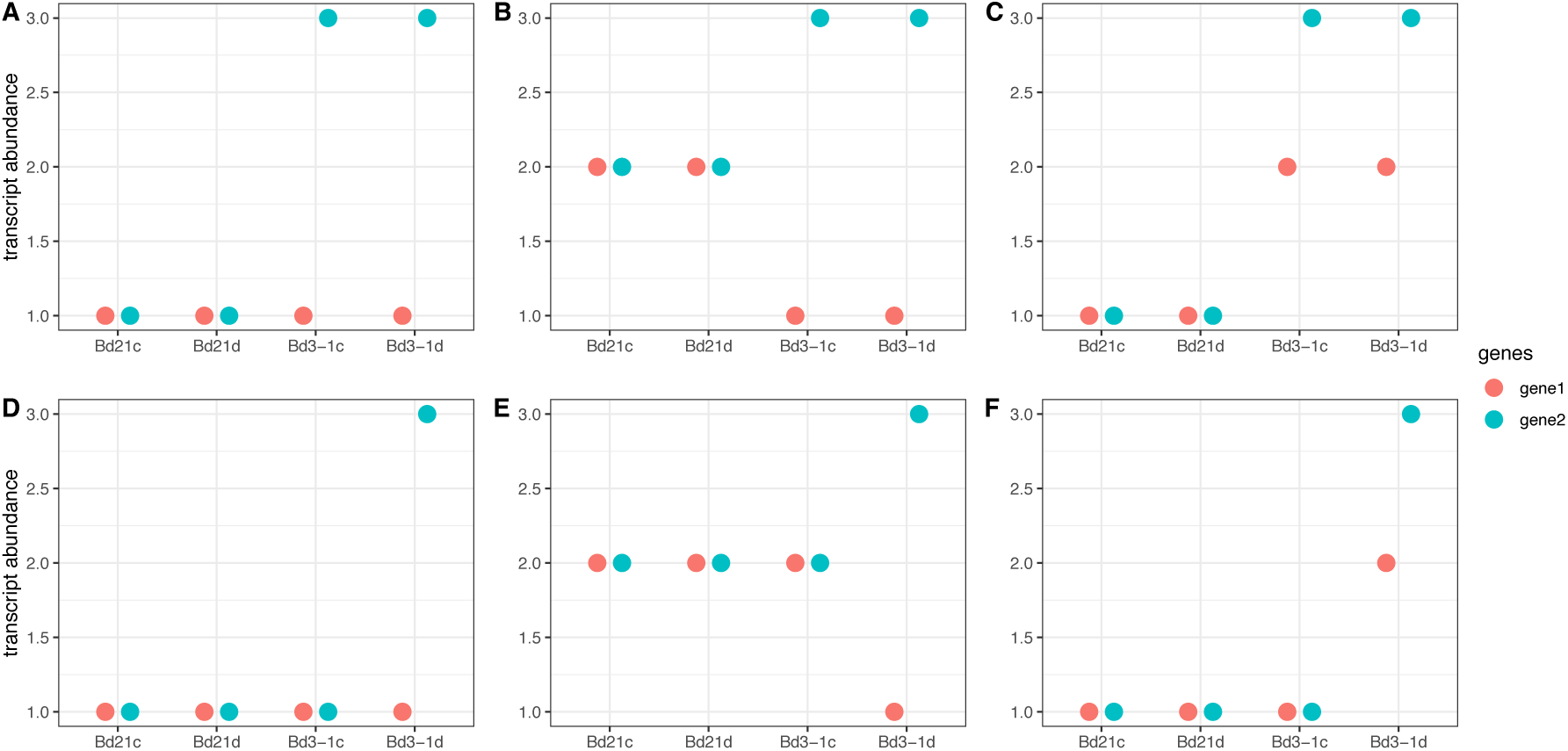

Examples illustrating gene-gene interaction. A-C) DGE G changes; E-G) DGE GxE changes. In each type of variation, the three plots show examples of different responses (A,D), direction (B,E), and sensitivity(C,F), respectively. Red and blue dots show two gene expression levels respectively, in four library types.

